# A Comparative Assessment of Human and Chimpanzee iPSC-derived Cardiomyocytes with Primary Heart Tissues

**DOI:** 10.1101/289942

**Authors:** Bryan J. Pavlovic, Lauren E. Blake, Julien Roux, Claudia Chavarria, Yoav Gilad

## Abstract

Comparative genomic studies in primates have the potential to reveal the genetic and mechanistic basis for human specific traits. These studies may also help us better understand inter-species phenotypic differences that are clinically relevant. Unfortunately, the obvious limitation on sample collection and experimentation in humans and non-human apes severely restrict our ability to perform dynamic comparative studies in primates. Induced pluripotent stem cells (iPSCs), and their corresponding differentiated cells, may provide a suitable alternative system for dynamic comparative studies. Yet, to effectively use iPSCs and differentiated cells for comparative studies, one must characterize the extent to which these systems faithfully represent biological processes in primary tissues. To do so, we compared gene expression data from primary adult heart tissue and iPSC-derived cardiomyocytes from multiple human and chimpanzee individuals. We determined that gene expression in cultured cardiomyocytes from both human and chimpanzee is most similar to that of adult hearts compared to other adult tissues. Using a comparative framework, we found that 50% of gene regulatory differences between human and chimpanzee hearts are also observed between species in cultured cardiomyocytes; conversely, inter-species regulatory differences seen in cardiomyocytes are found significantly more often in hearts than in other primary tissues. Our work provides a detailed description of the utility and limitation of differentiated cardiomyocytes as a system for comparative functional genomic studies in primates.

**Data availability and resource sharing:** Gene expression (RNA-seq) data are available at GEO under accession number GSE110471. All human and chimpanzee iPSCs are available upon request without restriction or limitation.

## Introduction

Comparative studies of humans and non-human apes are extremely restricted because we only have access to a few types of cell lines and to a limited collection of frozen tissues (Romero et al. 2012). In order to gain true insight into regulatory processes that underlie inter-species variation in complex phenotypes, we must have access to faithful model systems for a wide range of tissues and cell types. To address this challenge, we previously established a panel of iPSCs from human and chimpanzee fibroblasts (Gallego Romero et al. 2015; Burrows et al. 2016; Blake et al. 2017). We can use this comparative iPSC panels to derive multiple cell types representative of the three germ layers. For example, we recently differentiated the human and chimpanzee iPSCs into definitive endoderm cells (Blake et al. 2017) to study conservation in gene expression trajectories during early development. Our hope is that employing iPSC-based models from humans and chimpanzees will provide researchers with a dynamic and flexible system for comparative functional genomic studies in a large number of cell types.

Towards this goal, the purpose of the current study is to evaluate how well inter-species gene expression differences in heart are recapitulated in iPSC-derived cardiomyocytes. This effort is not unique; quite a few previous studies focused on characterizing similarities and differences between pluripotent stem cell derived cell types and their fetal and adult tissue counterparts in both human and mouse (Shinozawa et al. 2009; Roessler et al. 2014; Piccini et al. 2015; Uosaki et al. 2015; van den Berg et al. 2015; Handel et al. 2016; Li et al. 2016; Uosaki and Taguchi 2016; Xia et al. 2016; Bell et al. 2017). Generally, results from these studies have demonstrated that the derived cell types are most equivalent to fetal tissues rather than to the corresponding adult tissues. A few studies specifically explored protocol properties that may result in more mature derived cells (Nunes et al. 2013; Ogawa et al. 2013; Shan et al. 2013; Zhao et al. 2013; Uosaki et al. 2015; Ellen Kreipke et al. 2016; Youssef et al. 2016; Lin et al. 2017).

That said, these published works do not specifically address properties pertaining to the utility of differentiated cardiomyocytes in the context of a comparative study in primates. First, nearly all published studies were conducted using relatively few individuals (three or fewer), such that the observation of high similarity of gene expression patterns between cultured cells and primary tissue may be explained by lack of statistical power. Second, previous studies did not consider their observations in the broader context of other tissues or other species, so it is challenging to benchmark the observation of what is claimed to be ‘small or ‘large’ regulatory differences between tissues and cultured cells. Different protocols and batch effects make it difficult to perform meta-analysis of existing data to effectively address this issue. Finally, to date, no study that focused on the fidelity of differentiated cells included samples from chimpanzees.

To address these gaps, we performed a comparative study that was specifically designed to allow us to effectively compare gene expression data between cultured cardiomyocytes and primary hearts from humans and chimpanzees. Our study was also designed to allow us to benchmark the results against other primary tissues and across these two species. A key finding from previous studies pertaining to the fidelity of iPSC-derived cell types is the importance of cellular maturation after terminal differentiation. While the initial steps of cardiomyocyte differentiation are fairly well established, there remains some debate in the field as to the best method of cellular maturation (Veerman et al. 2015; Kolanowski et al. 2017). Common strategies include: treatment with triiodothyronine (T3) (Lee et al. 2010; Ivashchenko et al. 2013; Yang et al. 2014), electrical stimulation (Nunes et al. 2013) and temporal maturation in culture (Ivashchenko et al. 2013). The fidelity of cardiomyocytes to primary adult heart tissue is likely to increase with cellular maturation. Thus, to test the effects of temporal maturation and treatment with T3, we compared gene expression profiles from human and chimpanzee iPSC-derived cardiomyocytes at day 15, day 27 (with and without T3 treatment) post induction of differentiation with data generated from primary heart tissue from both species.

## Results

The primary goal of this work is not to test a particular hypothesis, but to develop an understanding of the degree to which iPSC derived cardiomyocytes can serve as a model system for comparative functional genomic studies in primates. To do so, we used matched panels of 9 human and 10 chimpanzee iPSC lines; a subset of these iPSCs was previously described (Gallego Romero et al. 2015; Burrows et al. 2016; Blake et al. 2017). Standard functional characterizations of iPSC lines, quality controls, and metadata for the entire panel of samples are provided in Supplemental Figures 1-3 and Supplemental Table 1.

**Figure 1.**
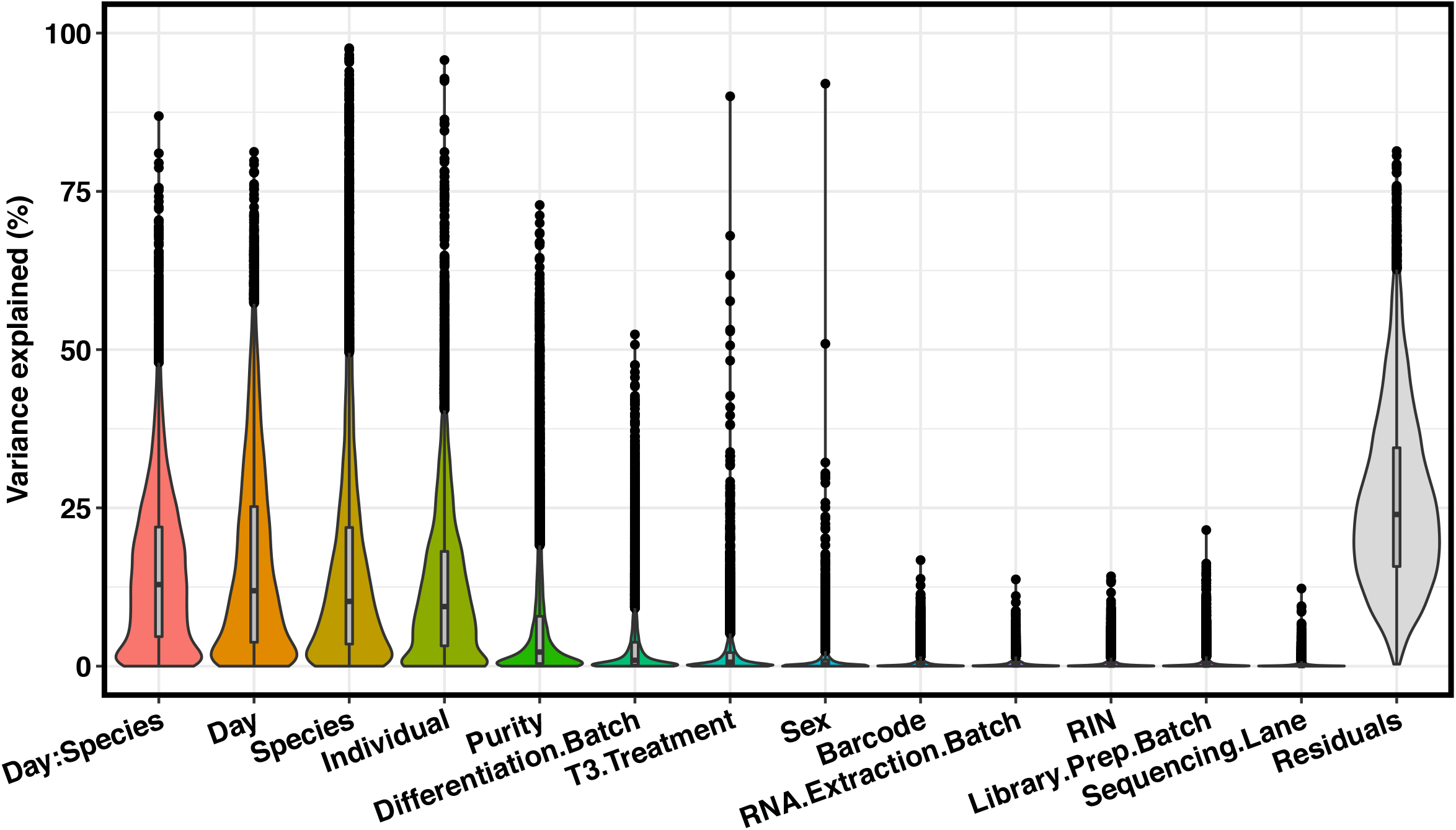
Variance partition. Results from variance partitioning analysis showing major drivers of gene expression variation in our study. All terms were modeled as fixed effects with the exception of individual, which was modeled as a random effect. “:” indicates day by species interaction term.

We differentiated the iPSC lines into cardiomyocytes using a recently developed protocol (Rana et al. 2012; Burridge et al. 2014; Burridge et al. 2015) with minor modifications (Methods). The same protocol was used for both the human and the chimpanzee iPSC lines. All of the cardiomyocyte cultures started beating 6-8 days post induction of differentiation. On days 20-27, a subset of cultures was treated with T3 (methods), which was shown to aid in the maturation of iPSC-derived cardiomyocytes (Lee et al. 2010; Yang et al. 2014; Lin et al. 2017). We assessed the purity of the iPSC-derived cardiomyocytes using flow cytometry (Methods) for cardiac troponin I (TNNI) and cardiac troponin T (TNNT2). By day 15, the purity of our samples was on par relative to the standard in the field (average purity of 68%; Supplemental Table 1). Importantly, we found no significant difference in sample purity across species (Supplemental Figure 4).

We performed the cardiomyocyte differentiations in two batches, with partial sample overlap across the batches; thus, our study contains technical replicates of differentiation for a subset of samples. In the first experimental batch, we collected RNA on day 27 from samples that were treated with T3. From the samples that were differentiated in the second batch, we collected RNA from the iPSCs, from differentiated cardiomyocytes on day 15, and again on day 27, from samples that were either treated or not treated with T3. To benchmark the gene expression data from the iPSC and iPSC-derived cardiomyocytes, we also collected RNA from post-mortem flash frozen heart tissues samples from 21 chimpanzees and 11 humans (Methods). Our overall study design (Supplemental Figure 4) affords us the flexibility to choose to analyze different subsets of samples that are balanced with respect to different technical and biological factors. We can thus effectively compare within and between species gene expression data from primary tissue and iPSC-derived cardiomyocytes that were harvested on days 15 and 27, with or without the treatment T3.

We examined and recorded the quality of the RNA samples from both tissues and cultures (Supplemental Table 1, Supplemental Figure 5), and sequenced each sample to an average coverage of 49.4 million total reads. We mapped the sequencing reads to an updated two-way human-chimpanzee orthologous exon reference set (Methods) and estimated relative gene expression levels using edgeR (Robinson et al. 2010). Throughout the differentiation and sample processing steps, we have recorded a large number of biological and technical properties (sample metadata available in Supplemental Table 1). As a first step of our analysis, we determined the potential impact of different properties of the study design and sample metadata on the observed gene expression levels (using variance partitioning; Methods). Most of the variation in our data can be explained by biological factors (species, day, and individual). Most study design properties explain only a modest amount of variation, but as expected, technical batch is associated with substantial amount of variation, as is sample purity (Figure 1). Importantly, neither sample purity nor technical batch are associated with species (Supplemental Figure 6).

### Which differentiated cardiomyocytes most resemble primary hearts?

Overall, we collected data from 110 samples (19 iPSCs, 59 differentiated cardiomyocytes, and 32 primary heart samples). To examine global trends in gene expression levels, we normalized data across all samples (using TMM; Methods), and visualized the data using principal component analysis (PCA; Figure 2A). We found that the major source of variation in the data is correlated with cell type; PC1 and PC2, which explain 36% and 22% of the variance, respectively, are highly associated with cell type (*P* < 10^-16^). The next major source of variation, as expected, is species (which is strongly associated with PC3; *P* < 10^-16^).

**Figure 2.**
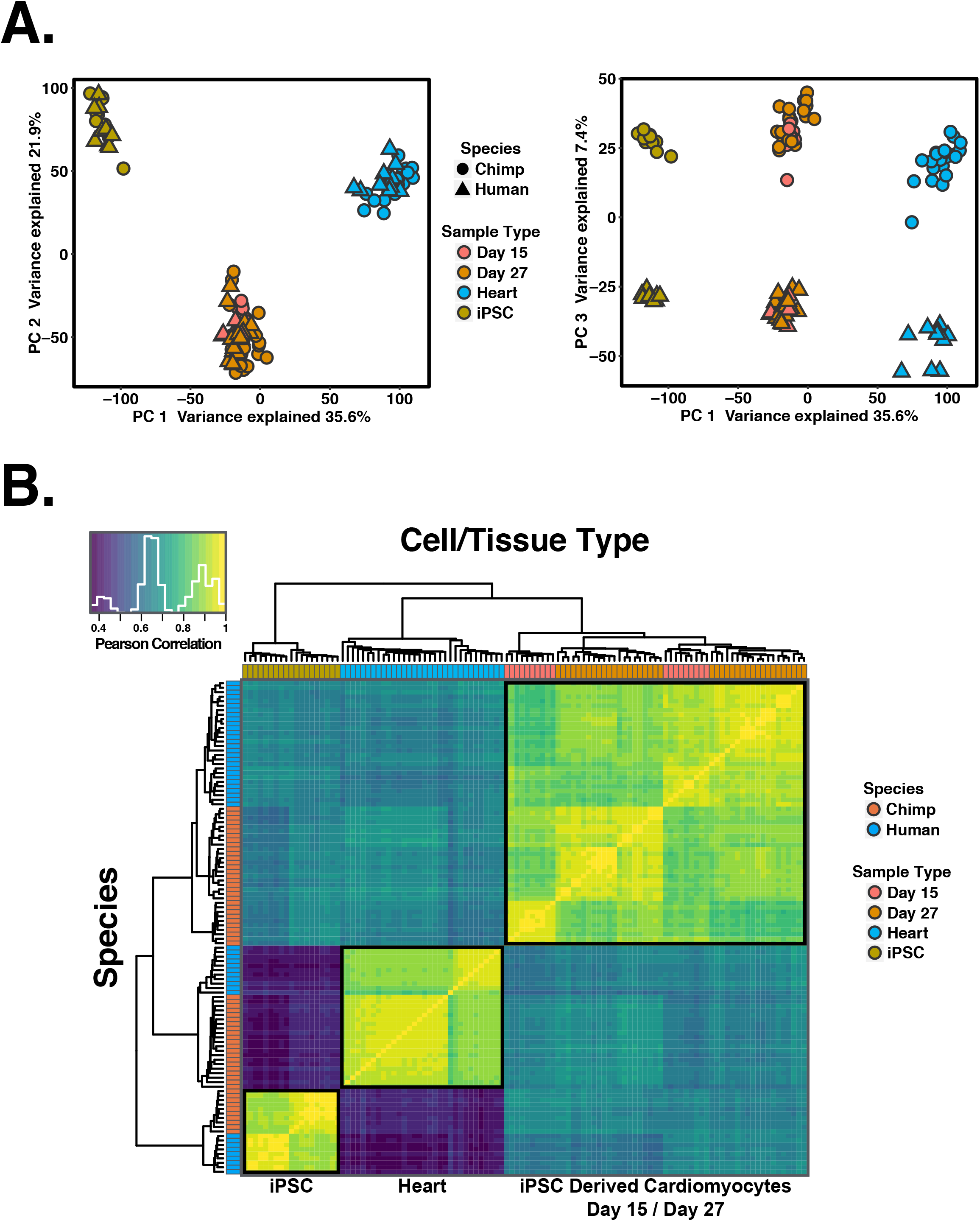
General patterns of gene expression variation. **A.** Normalized log2(RPKM) expression measurements for all genes projected onto the axes of the first two principal components. Color indicates cell or tissue type. Shape represents species. PC1 and PC2 are both strongly associated with cell or tissue type. **B.** Heat map of the pairwise Pearson’s correlation matrix of normalized log2(RPKM) gene expression values from 13,878 orthologous genes.

We performed unsupervised hierarchical agglomerative clustering on the correlation matrix of gene expression data, and found that the major trends seen in the PCA are recapitulated (Figure 2B). We noted that data from iPSC-derived cardiomyocytes clusters by species first, then by maturation day or treatment.

We focused on the differences between iPSC-derived cardiomyocytes at days 15 and 27, with and without T3 treatment. For this analysis, we considered within-species gene expression patterns. Using the data from each species independently, we asked which cardiomyocyte cultures are most similar to adult heart tissue with respect to their gene expression profiles. We first focused on temporal maturation (the untreated samples at days 15 and 27) and found that average pairwise correlation in gene expression data is higher between untreated day 27 cardiomyocytes and hearts than between day 15 cardiomyocytes and hearts, regardless of species (Figure 3A, paired t-test*; P* < 10^-16^).

**Figure 3.**
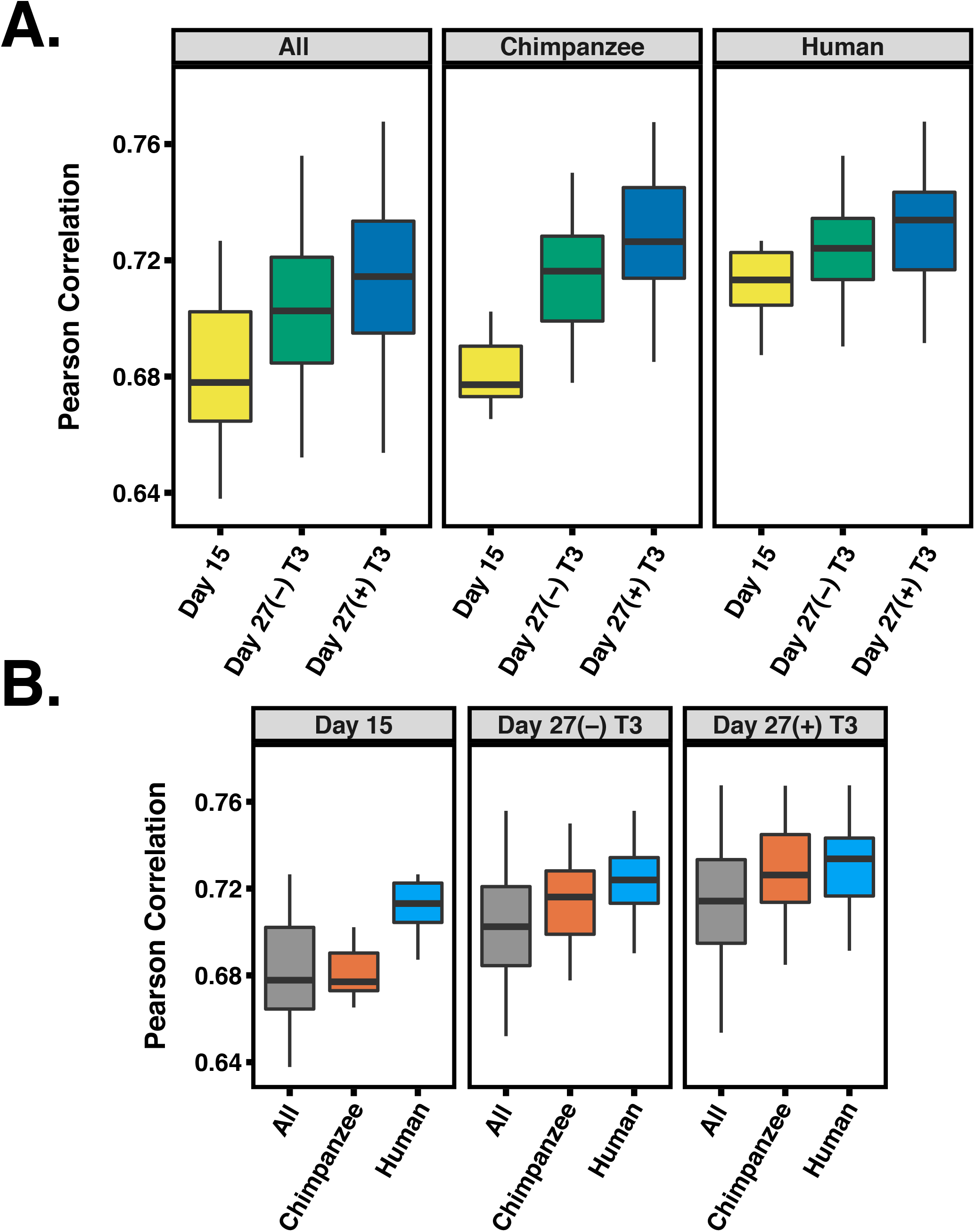
Pairwise correlations between heart tissue and different iPSC-derived cardiomyocytes. **A.** Pearson correlations between heart tissue, day 15, (-)T3 and (+)T3 day 27 iPSC-derived cardiomyocytes for chimpanzee, humans and All samples combined. **B.** Same data as in A, but reordered to show differences within a treatment across species.

This result indicates that temporal maturation increases the similarity between cardiomyocytes and adult heart tissue for both species. We proceeded to compare gene expression data between heart tissues and cardiomyocytes cultures at day 27 that were treated or not treated with T3. We found that gene expression data from samples treated with T3 are more similar to that of primary heart tissues, regardless of species (Figure 3A, *P* < 10^-4^). We compared the maturation effects across species and found that, at day 15, human cardiomyocytes are more similar to human adult heart tissue than chimpanzee cardiomyocytes are to chimpanzee adult hearts (Figure 3B, *P* < 10^-7^). However, by day 27, we did not observe any significant difference across species in this comparison (*P* = 0.13).

### Which genes are differentially regulated between hearts and cultured cardiomyocytes?

Based on our observations we concluded that as the cells mature, the fidelity of iPSC-derived cardiomyocytes as a model for primary hearts increases in both species. Moreover, T3 treatment helps to mature iPSC-derived cardiomyocytes, altering their gene expression such that it is more similar to that of adult tissues than temporal maturation alone. Yet, even T3 treated samples at day 27 are clearly distinct from heart samples (Figures 2 and 3). We thus characterized the specific regulatory difference between heart tissues and iPSC-derived cardiomyocytes.

To do so, we considered a subset of day 15 and T3 treated day 27 cardiomyocyte samples from 7 humans and 7 chimpanzees, all with purity above 50% (Supplemental Figure 7). We used a linear model framework to perform a combined analysis of all data from these samples, as well as from the corresponding iPSCs, and from the primary heart tissues (Methods). We focused on genes that were classified as differentially expressed between cell / tissue types in both humans and chimpanzees (Methods). At an FDR of 5% we classified 3,446 differentially expressed genes (of 13,878 orthologous genes) between samples at day 27 (T3 treated) and heart tissue, and 4,115 differentially expressed genes between samples at day 15 and heart tissue (Supplemental Table 2). For comparison, using the same approach, we classified nearly 8,000 differentially expressed genes between iPSCs and cardiomyocytes at either day 15 or day 27.

To gain insight into the processes that are differentially regulated between primary hearts and iPSC-derived cardiomyocytes, we considered GO functional annotations(Ashburner et al. 2000; The Gene Ontology 2017). We classified the differentially expressed genes between hearts and cardiomyocytes to (*i*) those that are not expressed at all in hearts, (*ii*) or in cardiomyocytes, and those that expressed in both tissues but are more highly expressed in (*iii*) heart or (*iv*) cardiomyocytes. Among genes that are expressed in hearts but not in cardiomyocytes, we found an almost exclusive enrichment for biological processes relating to immune responses and inflammation. In turn, among genes that are expressed in cardiomyocytes but not in hearts, we found enrichment of genes involved in cell cycle regulation. Among genes expressed in both cardiomyocytes and hearts, we found that genes involved in transcription regulation tend to be expressed at higher levels in cardiomyocytes and genes involved in metabolic processes specifically related to lipid metabolism tend to be expressed at higher levels in hearts (Figure 4; complete enrichment results in Supplemental Table 3; all reported enrichments associated with FDR < 0.05). Remarkably, when we consider all of the enrichment results with FDR < 0.05, we are able to account for 67% of genes that are classified as differentially expressed between hearts and iPSC-derived cardiomyocytes.

**Figure 4.**
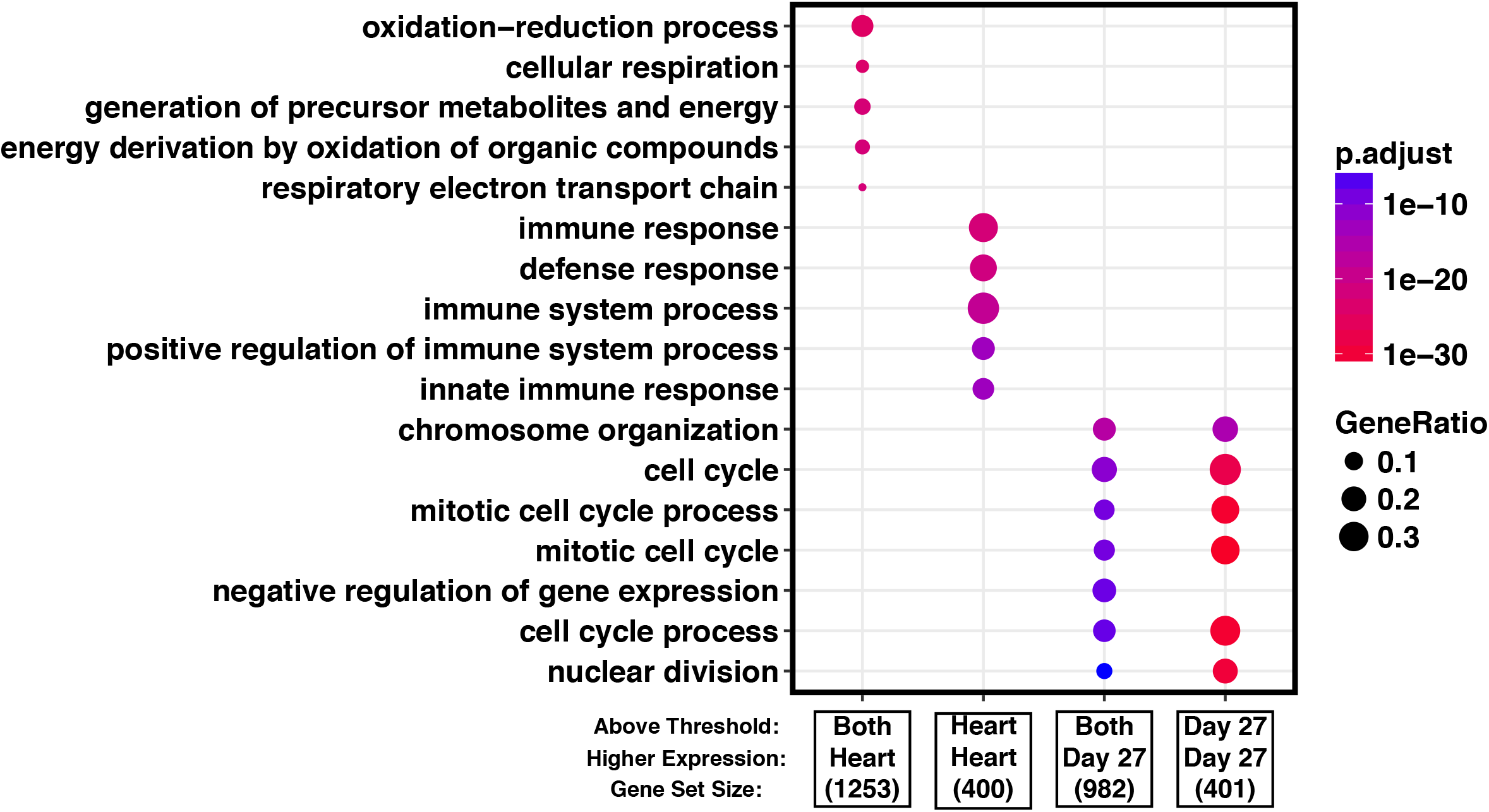
Gene enrichment analysis. We classified differentially expressed genes as those that are expressed at higher level (‘Higher Expression’) in either primary hearts or the iPSC-derived cardiomyocytes, and further, to those that are practically not expressed (‘Above Threshold’) in either primary hearts or the cardiomyocytes (see main text for more details). The top enrichments in GO biological processes for each of the categories is shown, gene list sizes are shown under the label for each gene set. Complete GO enrichment results are available in Supplemental Table 3. samples.

Up to this point, we considered gene expression patterns within species. To specifically examine iPSC-derived cardiomyocytes as a model system for comparative genomic studies, we proceeded to compare inter-species regulatory differences in either cell cultures or their corresponding primary tissues. We used a framework of linear modeling as above, but this time focused on the species by cell type interaction effect. To account for incomplete power when we classified genes as differentially expressed between species in any given cell or tissue type, we used a relaxed statistical cutoff for secondary observations. Namely, when a gene was classified as differentially expressed between species in one cell or tissue type using a stringent statistical cutoff (FDR of 5%), we used a relaxed statistical cutoff to classify the same gene as differentially expressed between species in the other cell or tissue type (nominal *P* < 0.05; Methods).

Using this approach, we found that roughly 50% of genes that are classified as differentially expressed between human and chimpanzee in heart tissues are also differentially expressed between cultured cardiomyocytes of the two species, regardless of whether we considered cardiomyocytes that were harvested at day 15 or day 27 (Figure 5). This result is robust with respect to the degree of purity of the cardiomyocyte cultures (within a range of 40-70%; Supplemental Figure 8, Methods) and statistical stringency (Supplemental Table 4). When we considered genes that are classified as differentially expressed between the species either in hearts or in cardiomyocytes (Methods), we found enrichment of similar functional terms when we considered either the day 27 cardiomyocytes or the heart tissue samples (Supplemental Table 5). Specifically, we found enrichment in terms relating to Carboxylic acid metabolism, cellular adhesion and xenobiotic metabolic processes. When we considered genes that are classified as differentially expressed between the species in both hearts and in cardiomyocytes, we found an enrichment of terms relating to multiple different metabolic processes, including monocarboxylic acid metabolic process, oxidation-reduction, and oxoacid metabolism (Supplemental Table 5, Supplemental Figure 9, all reported enrichments associated with FDR < 0.05).

**Figure 5.**
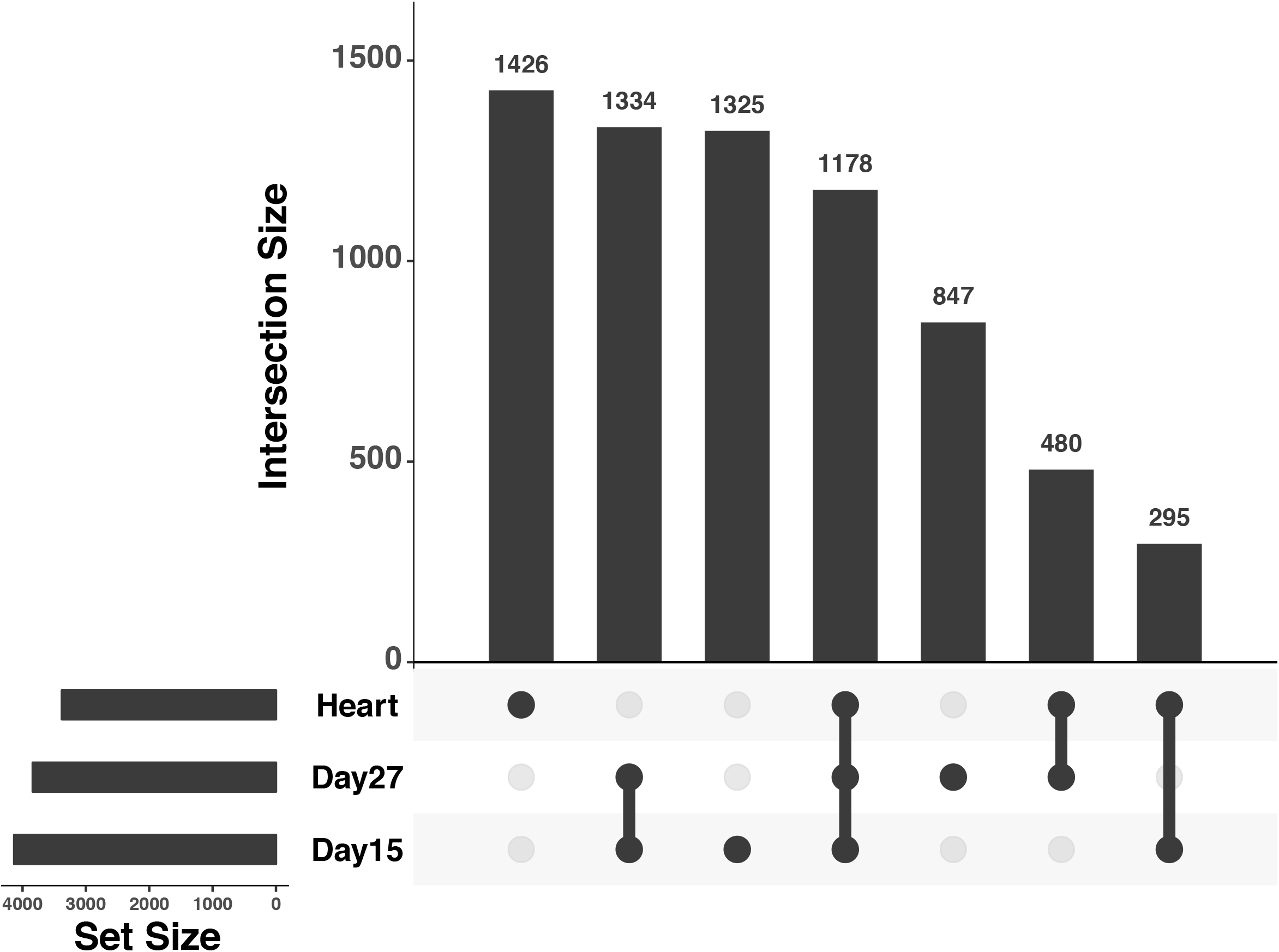
Upsetr diagram showing benchmarking results for human-chimpanzee DE genes using only iPSC-derived cardiomyocytes collected in this study. Total set sizes are shown on the bottom left, overlaps are shown by links with a filled circle, and the bars above the links shows corresponding size of a specific overlap of DE gene sets. To calculate the proportion of genes that are DE between human and chimpanzee in both heart and iPSC derived cardiomyocytes, overlaps between heart and the target cell type were summed together and divided by the total heart set size. Plot generated using R package Upsetr (Conway et al. 2017).

### Cardiomyocytes are more similar to hearts than to other primary tissues

Our observations thus far indicate that while much of the heart regulatory divergence between species can be captured by using iPSC-derived cardiomyocytes, there are also thousands of genes whose expression differs between hearts and cultured cardiomyocytes. To provide context for these observations, we used data from multiple publicly available RNA-seq studies, which include samples from four different adult tissues from both humans and chimpanzees, as well as from human fetal tissues (Bernstein et al. 2010; Brawand et al. 2011; Marchetto et al. 2013; Lin et al. 2014; Peng et al. 2015; Ruiz-Orera et al. 2015; Schultz et al. 2015; Uhlen et al. 2015; Williams et al. 2015; Wu et al. 2015; Barrette et al. 2016; He et al. 2016; Qin et al. 2016; Wang et al. 2016; Yan et al. 2016). We also included data from human and chimpanzee iPSCs, human ESCs, and human day 31 iPSC derived cardiomyocytes (Marchetto et al. 2013; Gallego Romero et al. 2015; Wu et al. 2015; Banovich et al. 2018). We processed all public data and our own newly collected data (from a total of 231 human samples and 111 chimpanzee samples) using a common analysis pipeline and a joint statistical model for the combined data (Methods).

We first focused on similarities and differences between tissues and cell types. We performed PCA using the combined data from all 342 samples, which suggested (based on visual inspection) separation of the samples by cell and tissue type in the first several principal components (Figure 6A; PC1 and PC2 are both highly associated with cell type or tissue type; *P* < 10^-16^). We only observed an association with species when we considered the loading on PC6 (*P* = < 10^-16^). We next performed unsupervised hierarchical agglomerative clustering using the correlation matrix of the combined gene expression data (Figure 6B). The samples clustered by cell and tissue type followed by species (with the exception of public data from one chimpanzee heart sample from (Peng et al. 2015)). Notably, we found that samples from iPSC-derived cardiomyocytes are consistently more similar to samples from fetal heart tissue, followed by samples from adult heart tissues than any other non-heart tissues (Figure 6B). These observations are consistent with previous findings that iPSC-derived cardiomyocytes are most similar to first trimester fetal hearts (van den Berg et al. 2015).

**Figure 6.**
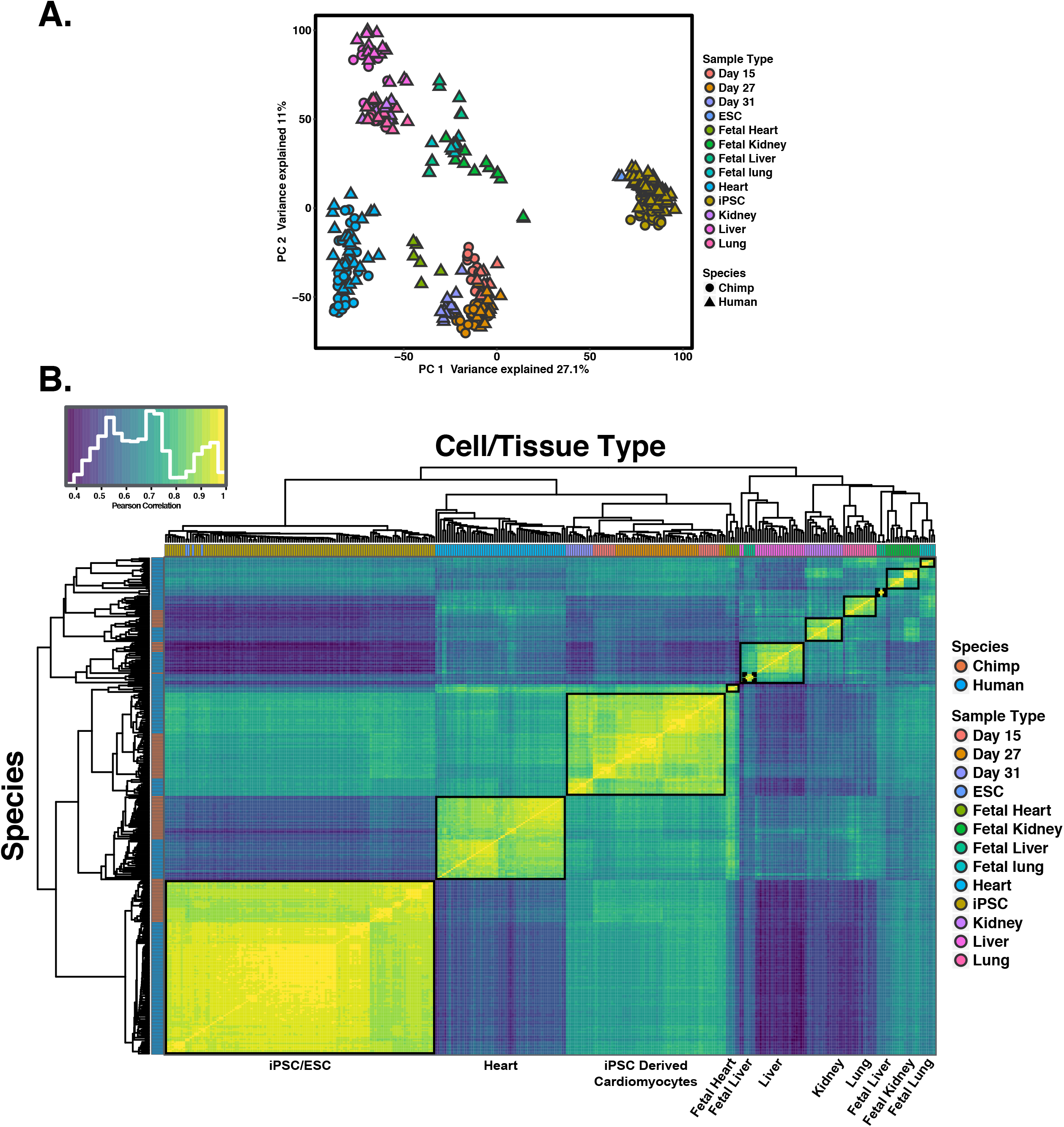
Global patterns of gene expression using multiple adult and fetal tissues from public datasets. **A.** Normalized log2(RPKM) expression measurements for all genes projected onto the axes of the first two principal components. Color indicates cell or tissue type. Shape represents species. PC1 and PC2 are both strongly associated with cell or tissue type. **B.** Heat map of the Pearson’s correlation matrix of normalized log2(RPKM) gene expression values from 335 samples for 15,829 orthologous genes. Each square represents the Pearson’s correlation of the normalized expression values between two samples.

We next evaluated the ability of iPSC-derived cardiomyocytes to capture regulatory divergence in hearts in the context of comparative data sets from other tissues. To do so, we used RNA-seq data collected from four individuals from each species from four different tissues: heart, lung, liver and kidney (GEO accession GSE112356). We classified genes as differentially expressed between human and chimpanzee in each tissue and in day 27 cardiomyocytes using a common analysis pipeline (Methods). To account for incomplete power when classifying genes as differentially expressed between human and chimpanzee in any given cell or tissue type, we again allowed for a more relaxed statistical cutoff for secondary observations. As expected, a large number of genes show inter-species differences in gene expression level across all tissues while different subsets of genes are differentially expressed between the species in only a single tissue (Supplemental Figure 10). Crucially, genes that are differentially expressed between human and chimpanzee in cardiomyocytes are much more likely to also be classified as differentially expressed between the species in hearts than in any other tissue (Figure 7).

**Figure 7.**
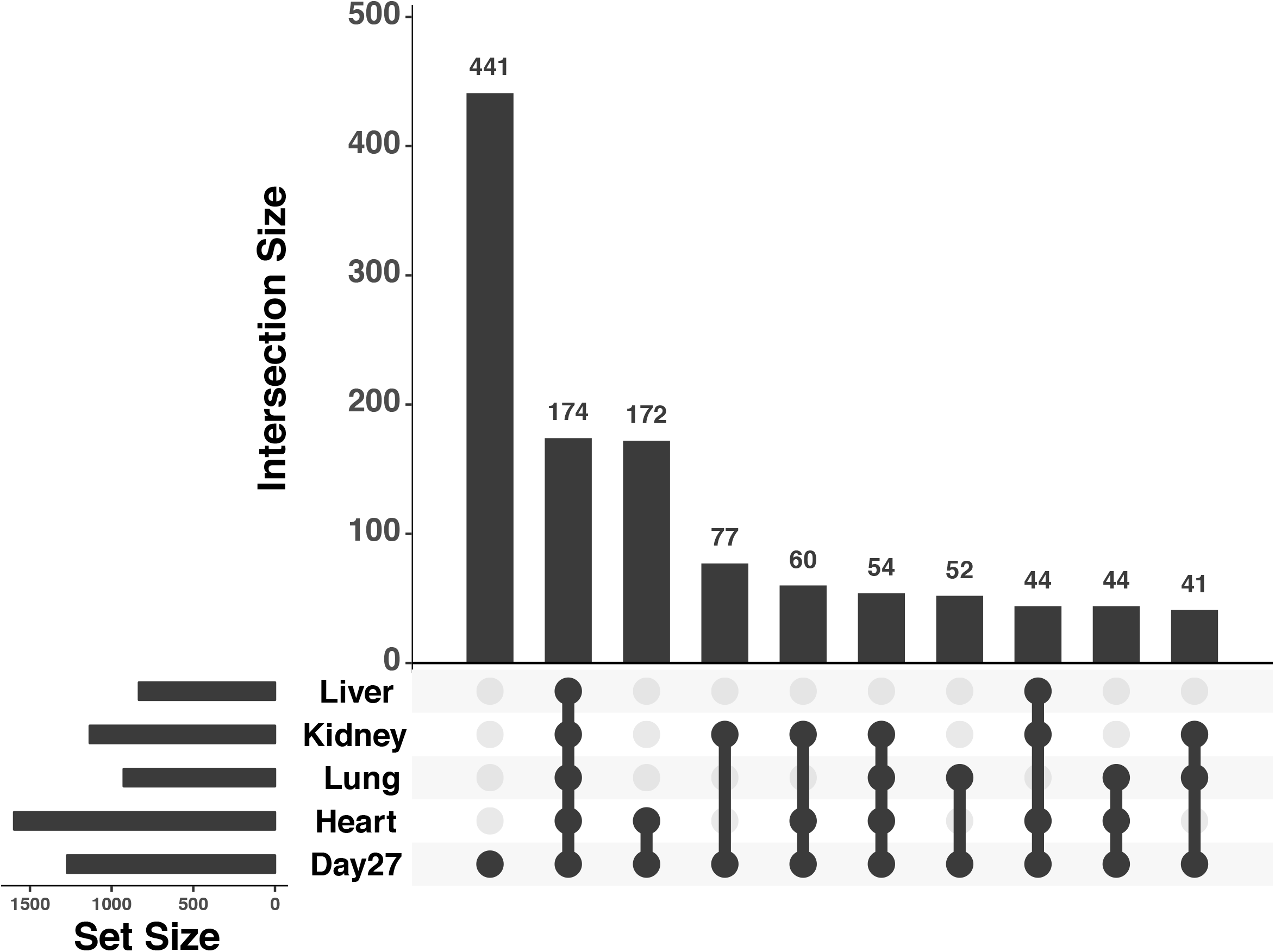
Upsetr diagram showing benchmarking results for iPSC-derived cardiomyocytes using independent tissue reference sets. Total set sizes are shown on the bottom left, overlaps are shown by links with a filled circle, and the bars above the links shows corresponding size of a specific overlap of DE gene sets. Plot generated using R package Upsetr (Conway et al. 2017).

## Discussion

The development of iPSC based model systems have the potential to transform the field of comparative primate genomics, assuming that differentiated cells can recapitulate biological processes that occur in primary cell types. Quite a few studies focused on gene regulatory variation in *in vitro* differentiated cells have already been published (including by our group), but a systematic assessment of the fidelity of these differentiated cells with respect to primary primate tissue has not been performed. The current study was therefore designed to specifically assess the degree to which differentiated cardiomyocytes can serve as a model system in which to study regulatory differences in human and chimpanzee hearts.

We debated which cell type to use for this first exploration of the fidelity of differentiated cells. Many different cell types are accessible using iPSC differentiation protocols (Murry and Keller 2008; Burridge et al. 2014; Loh et al. 2014; Clevers 2016). Despite the seemingly many options, in most cell types, individual variation still remains a current barrier to generate consistently pure cultures of target cell types (Osafune et al. 2008; Kattman et al. 2011; Nazareth et al. 2013; Burridge et al. 2015; Kyttala et al. 2016; Wernig et al. 2016). In contrast, cardiomyocyte differentiation protocols are relatively robust with respect to inter-individual variation (Burridge et al. 2015). This outcome is partially due to the dominance of a single pathway (WNT signaling) in resolving developmental bifurcations in favor of cardiomyocyte generation *in vitro* (Burridge et al. 2014; Rao et al. 2016). An additional advantage of utilizing a cardiac lineage for benchmarking is the relatively homogenous nature of cardiac muscle (e.g., the composition of the left ventricle is upwards of 40% cardiomyocytes (Zhou and Pu 2016)). With these factors in mind, differentiations of iPSCs to cardiomyocyte provide an optimal framework to begin characterizing the similarity of iPSC-derived cell types to primary tissue in primates.

We did not expect differentiated cardiomyocytes to be identical to hearts. That would not be a realistic expectation. We hoped that differentiated cardiomyocytes would be the most similar to hearts over than to any other tissue, and we hoped to shed light on the opportunities and limitations associated with the use of differentiated cardiomyocytes to study hearts. Our observations confirmed that gene expression data from differentiated cardiomyocytes are more similar to data from hearts than any other tissue we tested, but we also found thousands of differentially expressed genes between hearts and differentiated cardiomyocytes. The majority of these differentially expressed genes, however, may not point to inherent limitations of differentiated cardiomyocytes as a model, but rather they help us better understand the model properties.

Indeed, 67% of differentially expressed genes between hearts and differentiated cardiomyocytes may be explained, not by artifacts specific to the model, but by expected biological differences between *in vitro* cultured relatively pure cells and a complex primary adult tissue. While most myocyte populations in the adult heart no longer proliferate and have a restricted ability to reenter cell cycle (Foglia and Poss 2016; Mohamed et al. 2018), the differentiated cardiomyocytes, which resemble embryonic heart cells more than adult cells, likely retain the ability to reenter cell cycle similar to their fetal and preadolescent counterparts (Naqvi et al. 2014; Soonpaa et al. 2015). It thus may be expected that genes with functions related to transcription regulation and those involved in cell cycle regulation (there is a high overlap between these categories) would be more highly expressed in young differentiated cardiomyocyte cultures than in adult hearts. Additionally, the cultured cardiomyocytes are predominantly ventricular cardiomyocytes (Burridge et al. 2014), while the hearts contain a complex composition of cells, including atrial and ventricular cardiomyocytes, nodal cells, cardiac fibroblasts, endothelial cells, blood cells and resident immune cells. It is therefore reasonable that genes involved in immune responses and inflammation are highly expressed in hearts compared with differentiated cardiomyocytes. Finally, it has been previously reported that the metabolic state of hearts and cultured cardiomyocytes are different (Uosaki et al. 2015). Consistent with that report, we found that genes involved in lipid metabolism are highly expressed in hearts relative to differentiated cardiomyocytes, an expected observation given that our cell cultures were not supplemented with lipids. Thus, the majority of the thousands of differentially expressed gene between primary hearts and cardiomyocytes are not necessarily indicative of inherent limitations of the *in vitro* model. Rather, many of these regulatory differences point to biological (cell composition) and environmental (lipids) differences between cell culture and primary tissue.

When we specifically considered inter-species comparisons of gene expression levels using either cardiomyocytes or hearts, we found similar patterns. Namely, while most genes are classified as differentially expressed between humans and chimpanzee in both differentiated cardiomyocytes and hearts, thousands of genes are differentially expressed between the species in either hearts or cardiomyocytes. Yet, the discordant inter-species patterns in cardiomyocytes and hearts mostly involve genes with functions related to metabolic processes, regulation of gene expression, and cell cycle. As we discussed above, there are likely biological and environmental explanations to these patterns.

We thus conclude that iPSC-derived cardiomyocytes are a useful model with which to conduct comparative studies of gene regulation. We expect that iPSC-derived cardiomyocytes will be mainly used to comparatively study dynamic gene regulatory processes, such as during differentiation, when frozen heart samples – even when available – are unsuitable. The model is not perfect, but our study indicates that regulatory patterns observed in cultured cardiomyocytes cannot be mistaken as anything other than a representation of the biological processes that occur in primary hearts. With that in mind, we recommend that the data from our study, which are available in unprocessed and processed forms, should be used to evaluate the likelihood that future observations based on cultured cardiomyocytes faithfully represent regulatory patterns in primary heart tissues.

## Methods

### iPSC panel

In this study, we include 8 chimpanzee iPSC lines and 6 human lines from previously described panels (Gallego Romero et al. 2015; Burrows et al. 2016; Blake et al. 2017). We also include 2 chimpanzee iPSC lines and 3 human lines not previously described (characterizations for all lines used in the paper are provided in Supplemental Figures 1-3). All iPSC lines are matched for cell type of origin, reprogramming method, and culture conditions. Original chimpanzee fibroblast samples for generation of iPSC lines were obtained from the Yerkes Primate Center under protocol 006–12. Human fibroblasts samples for generation of iPSC lines were collected under University of Chicago IRB protocol 11–0524. Feeder free iPSC cultures were initially maintained on Growth Factor Reduced Matrigel using Essential 8 Medium (E8) as previously described (Gallego Romero et al. 2015). After 10 passages in E8, all cell lines were transitioned to a 50/50% ratio of iDEAL/E8 feeder free medium that was prepared in house as specified previously (Marinho et al. 2015). Cell culture was conducted at 37°C, 5% CO_2_, and atmospheric O_2_.

### Cardiomyocyte differentiation

Cardiomyocytes were generated following recently published protocols (Rana et al. 2012; Burridge et al. 2014; Burridge et al. 2015) with minimal modification. At 12 hours prior to initiating differentiation, iPSC lines at 70-90% confluence were seeded to achieve a starting density of 150,000-250,000 cells/cm^2^. Differentiations were initiated by removing stem cell maintenance media and adding RPMI base media (with HEPES LifeTechnologies# 22400105) supplemented with 6 μM CHIR9902 (LClabs# C-6556), 0.5 mg/mL BSA (Sigma-Aldrich# A2153), 0.213 mg/mL ascorbic acid (Santa Cruz Bio# sc-228390), 0.3 μg/mL sodium selenite (Sigma-Aldrich# S5261), 5 μg/mL Holo-transferrin (Sigma-Aldrich# T3705), 1 μg/mL Linolenic acid (Sigma-Aldrich# L2376), 1 μg/mL Linoleic acid (Sigma-Aldrich# 1012) and 1x Pen/Strep (Days 0-1 media). After 48 hours, the media was changed to RPMI base media (with HEPES LifeTechnologies# 22400105) supplemented with 2 μM Wnt-c59 (Tocris# 5148), 0.5 mg/mL BSA (Sigma-Aldrich# A2153), 0.213 mg/mL ascorbic acid (Santa Cruz Bio# sc-228390 and 1x Pen/Strep (Days 2-3 media). After another 48 hours, the media was changed to RPMI base (with HEPES LifeTechnologies# 22400105) supplemented 0.5 mg/mL BSA (Sigma-Aldrich# A2153), 0.213 mg/mL ascorbic acid (Santa Cruz Bio# sc-228390) and 1x Pen/Strep. Media was changed every 48 hours until day 10 (total of 2 additional media changes). At day 10 the media was switched to RPMI media (no glucose, Cellgro#10-043-CV) supplemented with 5 mM Sodium D/L Lactate (Sigma-Aldrich# L4263), 0.5 mg/mL BSA (Sigma-Aldrich# A2153), 0.213 mg/mL ascorbic acid (Santa Cruz Bio# sc-228390) and 1x Pen/Strep, media was changed every 48 hours (total of 2 additional media changes). At day 15, cells were dissociated using TrypLE and split to a density of 0.4 x 10^6^ cells/cm^2^ onto Matrigel coated dishes. Cells were replated in RPMI media (no glucose, Cellgro#10-043-CV) supplemented with 5% FBS, 10 mM D-galactose (Sigma-Aldrich# G5388), 1 mM Sodium Pyruvate (LifeTechnologies# 11360070), 1X NEAA (LifeTechnologies# 11140076), 5 mM HEPES (LifeTechnologies# 15630080), mM Sodium D/L Lactate (Sigma-Aldrich# L4263), 0.5 mg/mL BSA (Sigma-Aldrich# A2153), 0.213 mg/mL ascorbic acid (Santa Cruz Bio# sc-228390) and 1x Pen/Strep (Galactose media). After 24 hours, the media was changed to Galactose media without FBS. Media was changed every 48 hours (total of 1 additional media change). At day 20 media was changed to Galactose media supplemented with 3 ng/mL Triiodothyronine (T3, Sigma-Aldrich# T6397) for T3(+) cardiomyocyte cultures. Media was changed every 48 hours until cells were harvested at day 27 (total of 3 additional media changes). For the first two media changes (Days 0 and 2) cultures were supplied with an excess of media (0.499 mL/cm^2^), all other days’ media was added at (0.2495 mL/cm^2^).

### Determining iPSC-derived cardiomyocyte purity using flow cytometry

Day 15 cells were dissociated as described above, and aliquots were pulled off for flow analysis prior to replating. Day 27 iPSC-derived cardiomyocytes were dissociated by 10 minute incubations with TrypLE supplemented with 0.5 U/ml Liberase TH. Dissociated cells were centrifuged at 200 x g for 5 minutes at 4°C and washed with PBS. Cells were fixed and stained as specified previously (Burridge et al. 2014; Burridge et al. 2015). Briefly, 0.5-1 million cells were fixed using a 1% PFA solution. Cells were fixed at 4°C for 30 minutes before washing once using FACS buffer (autoMACS Running Buffer, Miltenyi Biotech). Fixed cells were permeabilized by incubating with cold 90% methanol for 30 minutes at 4°C. Fixed, permeabilized cells were washed 2x using FACS buffer prior to staining. For immunostaining, 150,000 cells were transferred to BRAND lipoGrade 96 well immunostaining plates and centrifuged at 200 x g for 5 minutes at 4°C. Cells were rinsed in FACS buffer then resuspended in the staining solution. A single master mix containing 0.5% BSA/0.1% Triton X-100, PE-labeled TNNT2 (Bdbio clone 13-11) and Alexa Flour 647 labeled-TNNI3 (BDbio clone C5) All antibodies were used at the manufacturer recommended dilution. Cells were stained for 1 hour at 4°C and subsequently washed 3x with a PBS solution containing 0.5% BSA/0.1% Triton X-100. Stained cells were resuspended after the final spin in 150 μL FACS buffer containing 5 μM Vybrant DyeCycle Violet Stain (Thermofisher #V35003) for acquisition on a BD LSR II flow cytometer. After data acquisition, we used the program FlowJo (https://www.flowjo.com/) to determine compensation scaling. To do so, we used data from single stained compensation beads (Life Technologies) that were stained and collected in parallel. Live, intact, single cells were gated based on FSC and SSC channels. To determine gating for dual positive iPSC-derived cardiomyocytes, we used a biological negative (an iPSC line stained in parallel) to determine appropriate positive signal for gating.

### Isolation of RNA and DNA

To isolate RNA and DNA from cardiomyocytes and iPSC lines, we first aspirated old cell culture media, then rinsed culture wells using PBS. After aspirating PBS, 300 μL of DNA/RNA lysis buffer was added directly to culture plates to lyse cells on the plate. The entire lysate was transferred to a 1 mL tube and immediately frozen for extraction with other samples in balanced batches. We extracted the RNA using the ZR-Duet DNA/RNA MiniPrep kit (Zymo) with the addition of an on column DNAse I treatment step prior to RNA elution.

Post-mortem human heart tissues were provided by the National Disease Research Interchange (NDRI). Chimpanzee post-mortem heart tissues were provided by Yerkes primate center, the Southwest Foundation for Biomedical Research, and the New Iberia Research Center, MD Anderson Cancer Center. For lysis of heart tissue, frozen tissue from the left ventricle was broken down into smaller chunks while working on dry ice to prevent thawing, and subsequently weighed in 1.5 mL tubes to determine weight of tissue. For each mg of tissue, we added 8 μL of 1X RNA/DNA shield buffer. Tissues were homogenized in RNA/DNA shield using sterile plastic pestles. To further homogenize tissues and release RNA and DNA, tissues were digested with Proteinase K for 30 minutes at 55°C. Digested, homogenized tissues were centrifuged for 2 minutes to pellet debris, the volume of supernatant was measured and was transferred to fresh tubes. An equal volume of DNA/RNA lysis buffer was added and the solution was mixed prior to freezing at −80. Frozen samples were thawed and extracted in parallel using the same protocol as used for cells.

### Sequencing library preparation and RNA sequencing

We used non-strand specific, polyA capture to generate RNA-seq libraries according to the Illumina TruSeq protocol. To estimate the RNA concentration and quality, we used the Agilent 2100 Bioanalyzer. We added barcoded adaptors (Illumina TruSeq RNA Sample Preparation Kit v2) and sequenced the 100 base pair single-end RNA-seq libraries on the Illumina HiSeq 4000 at the Functional Genomics Core at University of Chicago on two flow cells. We used FastQC (http://www.bioinformatics.babraham.ac.uk/projects/fastqc/) to confirm that the reads were high quality.

### Mapping of RNA-seq data, orthologous exons, read count transformation and normalization

We mapped human reads to the hg38 genome and chimpanzee reads to panTro5 using HISAT2 (Kim et al. 2015) using the default parameters. We kept on only reads that mapped uniquely. To prevent biases in expression level estimates due to differences in mRNA transcript size and the relatively poor annotation of the chimpanzee genome and differences in mappability between species, we only kept reads that mapped to a list of orthologous metaexons. As the previous version of this metaexon database was generated using older versions of the genomes, we generated an updated orthoexon database based on the most current genome builds (see methods below for details). Gene expression levels were quantified using the FeatureCounts function in SubRead (version 1.5.1, (Liao et al. 2014) We performed all downstream processing and analysis steps in R unless otherwise stated.

We used a common analysis pipeline to process RNA sequencing data collected from the current study and also from the downloaded public datasets. We transformed raw gene counts into log_2_-transformed counts per million (CPM) using edgeR (Robinson et al. 2010). To filter for the lowly expressed genes, we kept only genes with an average expression level of log_2_(CPM) > 2 in at least one cell type (Robinson and Oshlack 2010). For the remaining genes, we normalized the read counts using the weighted trimmed mean of M-values algorithm (TMM) (Robinson and Oshlack 2010) to account for between-sample differences in the read counts at the extremes of the distribution and calculated the TMM-normalized log_2_-transformed CPM. After TMM-normalization we then performed a cyclic loess normalization with the function normalizeCyclicLoess from the R/Bioconductor package limma (Ballman et al. 2004; Ritchie et al. 2015). To account for gene length differences between species we calculated normalized log_2_-transformed RPKM values by using the function rpkm with normalized library sizes from the package edgeR (Robinson et al. 2010). We measured the “gene lengths” as the sum of the lengths of the orthologous exons.

To estimate the total gene expression variation attributed to the different properties of study design and sample metadata in our study, we used a linear mixed model implemented in variancePartition (Hoffman and Schadt 2016). We modeled all effects as fixed effects with the exception of individual which was modeled as a random effect.

### Updating orthologous exons

As part of the work contained in this paper, we updated our previously generated exon database (Blekhman 2012) with slight modification. We describe the modifications as follows. First, exon definitions were initially obtained from Ensembl release 88 (downloaded March 2017 containing 744,360 exons across 63,898 genes). This initial set of exon definitions was condensed to remove redundancies (haplotypes, patches and alternate exon entries resulting from different source annotations) and exons from genes that were fully overlapping. Regions of exons containing overlaps belonging to different genes were removed; exons with at least 10 bp of sequence not contained within the overlapping region were retained by removing only the region of overlap. This resulted in the division up of some exons into one or more smaller unique exons. Exons for the same genes with overlaps of at least 1 bp were collapsed into single metaexons containing all overlapping exons. Human fasta sequences (for the reduced set of 320,753 meta exons across 56,479 genes) were then extracted from the genome (Ensembl GRCh38.p10). We then used BLAT (BLAT V. 35 (Kent 2002)) to identify the orthologous exon sequences in the chimpanzee genome (panTro5). All hits with indels larger than 25 bp were removed (using a function blatOutIndelIdent, from https://bitbucket.org/ee_reh_neh/orthoexon), the fasta sequences for the hits with the highest sequence identity were extracted from the genome (using Bedtools (Quinlan and Hall 2010)). The chimpanzee sequences were then blatted against the human genome and back to the chimpanzee genome. Queries that did not return the original blat locations (for either chimpanzee or human) were removed, as were entries where multiple possible mappings occurred with higher than 90% sequence identity. Finally, different exons mapping to the same location (different locations in human, but same location in chimpanzee) were removed. This updated human-chimpanzee orthologous exon database resulted in 254,172 metaexons across 44,125 genes and is provided as Supplemental File 1.

### Linear modeling, differential expression and gene enrichment analyses

Differential expression analysis was performed using an empirical Bayes approach to linear modeling in genome wide expression data implemented in the R package limma (Smyth 2004; Smyth et al. 2005). To apply this linear modeling approach to RNA-seq read counts, we calculated weights that account for the mean-variance relationship of the count data using the function voom from the limma package (Law et al. 2014).

In analyses with both tissue and iPSC-derived cardiomyocytes, we cannot correct for purity differences, thus, we chose iPSC-derived cardiomyocytes with the highest purity (Supplemental Table 1). Since RNA quality varied greatly between tissues and derived cells (Supplemental Table 1, Supplemental Figure 5), we modeled RNA quality as a fixed effect using RIN scores. For all pairwise differential expression comparisons, the species, cell type, RNA quality (RIN), and a species-by-cell type interaction were modeled as fixed effects, and individual as a random effect. We used contrast tests in limma to find genes that were differentially expressed between tissues and cells for each species. For each pairwise differentially expressed (DE) test, we corrected for multiple testing with the Benjamini & Hochberg false discovery rate (Benjamini and Hochberg 1995) and genes with an FDR-adjusted *P* values < 0.05 were considered DE.

To identify genes as shared or specific to one species when examining DE genes between different cell or tissue types while accounting for incomplete power to detect overlaps, we first identified DE genes between the different cell or tissue types at an FDR of 5% in each species separately. Using this list of significant DE genes, we then looked in the other species, but classified genes as DE with a more relaxed cut off using a nominal *P* value of 0.05. Genes identified at an FDR of 5% in one species but not significant at a nominal *P* value of less than 0.05 in the other species were determined to be specific to one species. Genes that were significant in one species at FDR of 5% and significant at a nominal *P* value of less than 0.05 in the other species were designated as shared. This same approach was also taken when looking at interspecies DE genes in heart tissue or iPSC-derived cardiomyocytes.

When examining the overlap of interspecies DE genes in multiple different tissues, data for each tissue or cell type was modeled separately, as all data could not be modeled together due to differences in sequencing technology (50 bp Illumina GAIIx vs 100 bp Illumina HiSeq 4000). For this analysis, we only included terms for species and RNA quality (RIN score). To identify shared interspecies DE genes across different tissues and cell types while accounting for incomplete power to detect overlaps, we used the same approach as detailed above.

To identify significantly enriched GO biological process terms for genes DE between day 27 cardiomyocytes and heart tissue, we used the compareCluster function implemented in the R package clusterProfiler (version 3.4.4, (Yu et al. 2012)). We used all expressed genes as our background for enrichment and only considered GO terms containing more than three annotated genes and less than 3000 genes. When considering all interspecies differentially expressed genes in heart tissues or day 27 samples, we found no significantly enriched GO terms. Results in the main text were derived from GO analysis using genes with an absolute log_2_-fold change between human and chimpanzee higher than 1.5.

### Determining how purity differences affect estimates of interspecies DE gene overlaps

Since target cell type heterogeneity may vary within tissues and cannot be easily assessed, we were interested in the effect of purity on our ability to detect interspecies tissue specific DE patterns using iPSC-derived cardiomyocytes. As we are unable to physically alter (or effectively estimate) the composition of post mortem tissues, we instead used different combinations of T3 treated day 27 cardiomyocyte samples to generate 277 different average purity scenarios (Supplemental Figure 11). With each combination of samples, we repeated the same analysis as described in the main text to determine how many interspecies DE genes identified in heart tissue could be recapitulated using iPSC-derived cardiomyocytes.

## Acknowledgment

We thank Michelle Ward, Joyce Hsiao and other members of the Gilad lab for helpful discussions and comments on the manuscript. This work was supported by NIH grants GM120167, HL139447 as well as the Yerkes National Primate Research Center Base Grant ORIP/OD P51OD011132. BJP was supported by the Training in Emerging Multidisciplinary Approaches to Mental Health and Disease (T32MH020065). LEB was supported by the National Science Foundation Graduate Research Fellowship (DGE-1144082) and by the Genetics and Gene Regulation Training Grant (T32 GM07197). JR was supported by the Marie Curie fellowship (IOF 273290). The content presented in this article is solely the responsibility of the authors and the funding bodies had no role in the design of the study and collection, analysis, interpretation of data, and in writing the manuscript.

---

**Supplemental Figure 1.**
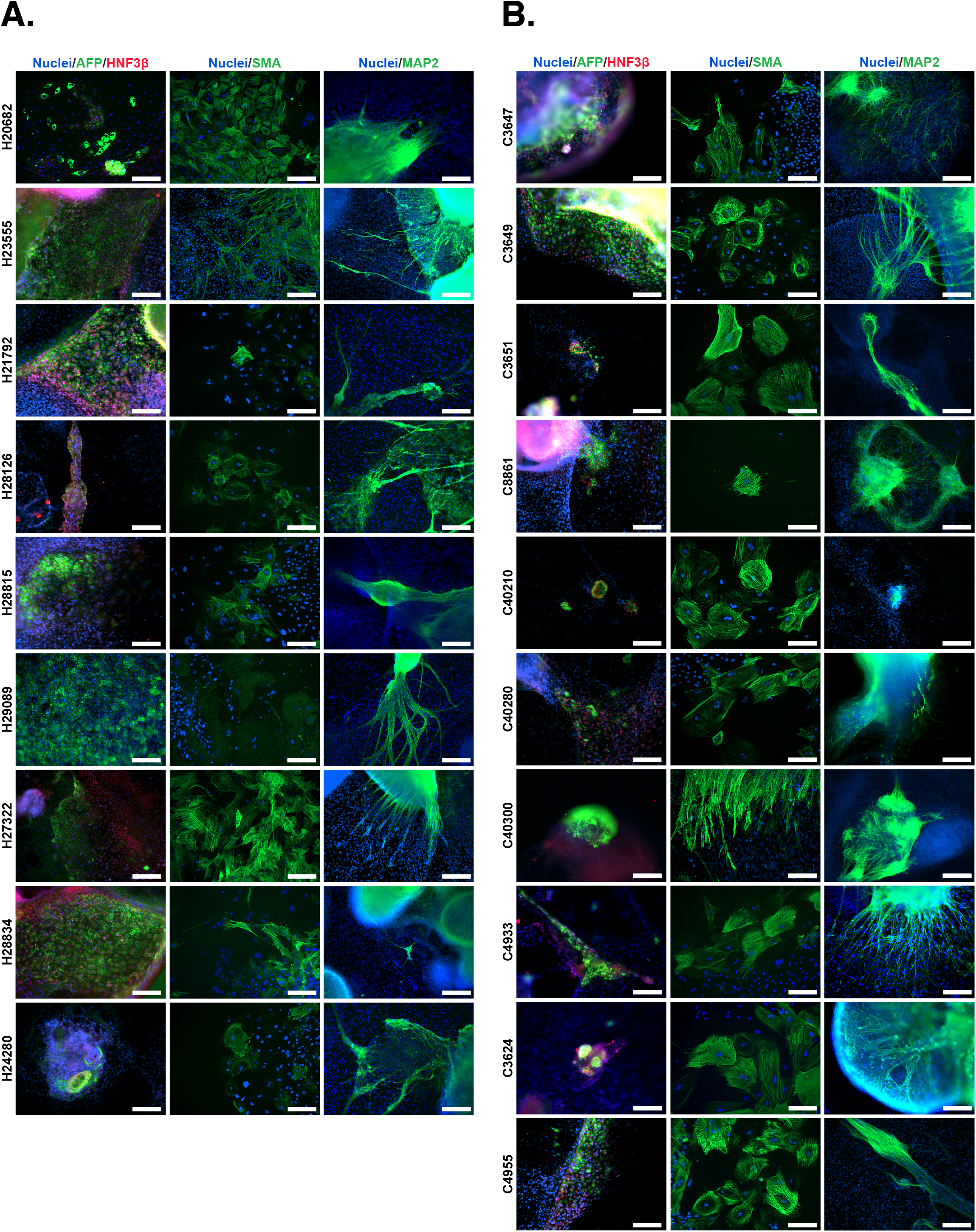
Spontaneous differentiation assay of pluripotency for iPSC lines used in this study. **A.** Immunocytochemistry (ICC) staining of spontaneously differentiated embryoid bodies for Human iPSC lines. **B.** Immunocytochemistry (ICC) staining of spontaneously differentiated embryoid bodies for chimpanzee iPSC lines, antibodies identifying cell types derived from the three germ layers as indicated. Scale bar: 200 μM.

**Supplemental Figure 2.**
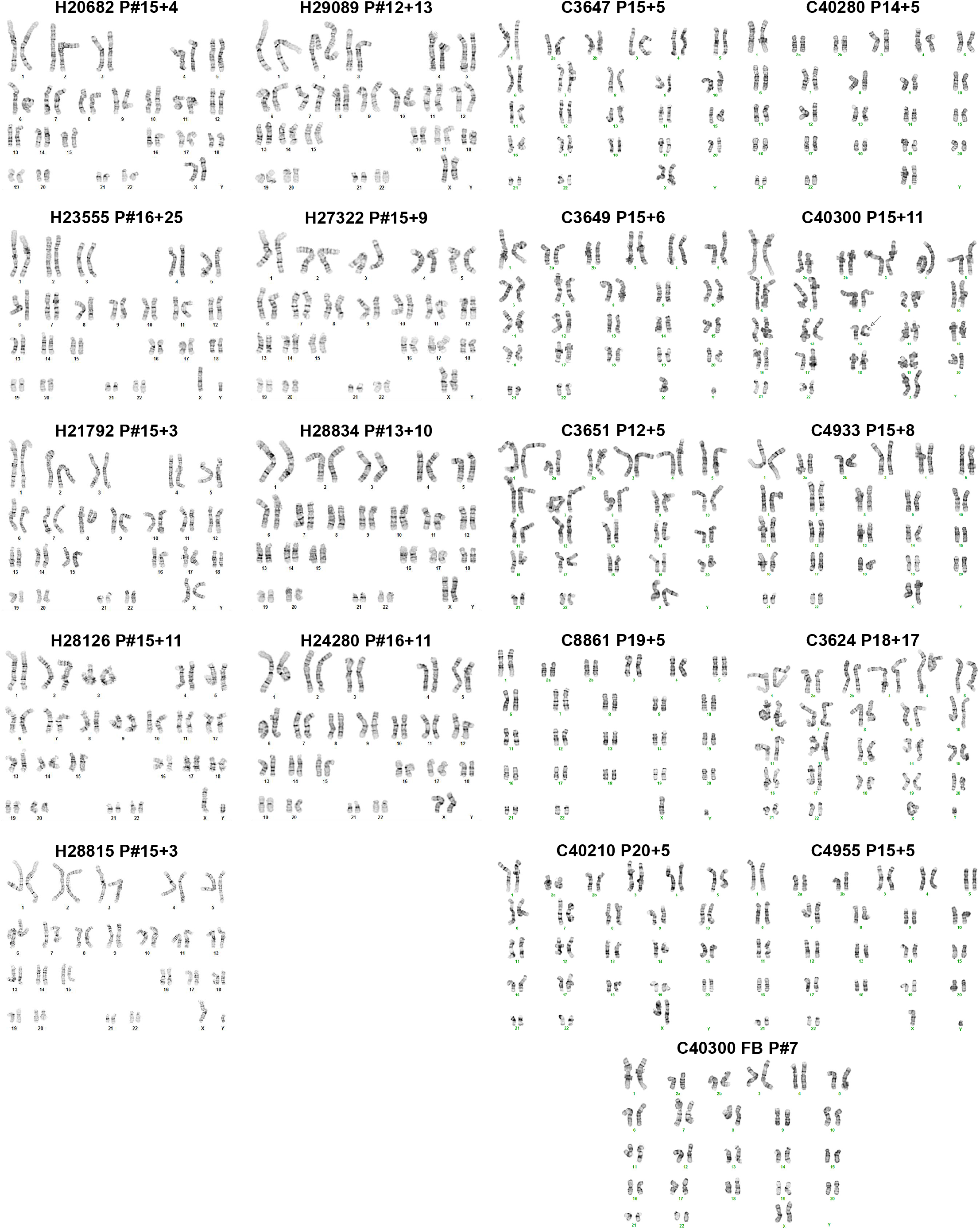
Karyotypes for human and chimpanzee iPSC lines. Karyotypes for human (right) and chimpanzee lines (left) used in this study. We identified additional bands in the p-arms of one chromosome 13 homolog and one chromosome 18 homolog for chimpanzee iPSC line C3. Thus, we tested the source fibroblast line (C3 FB) to determine that these polymorphisms were normal polymorphisms and not formed de novo as a result of reprogramming.

**Supplemental Figure 3.**
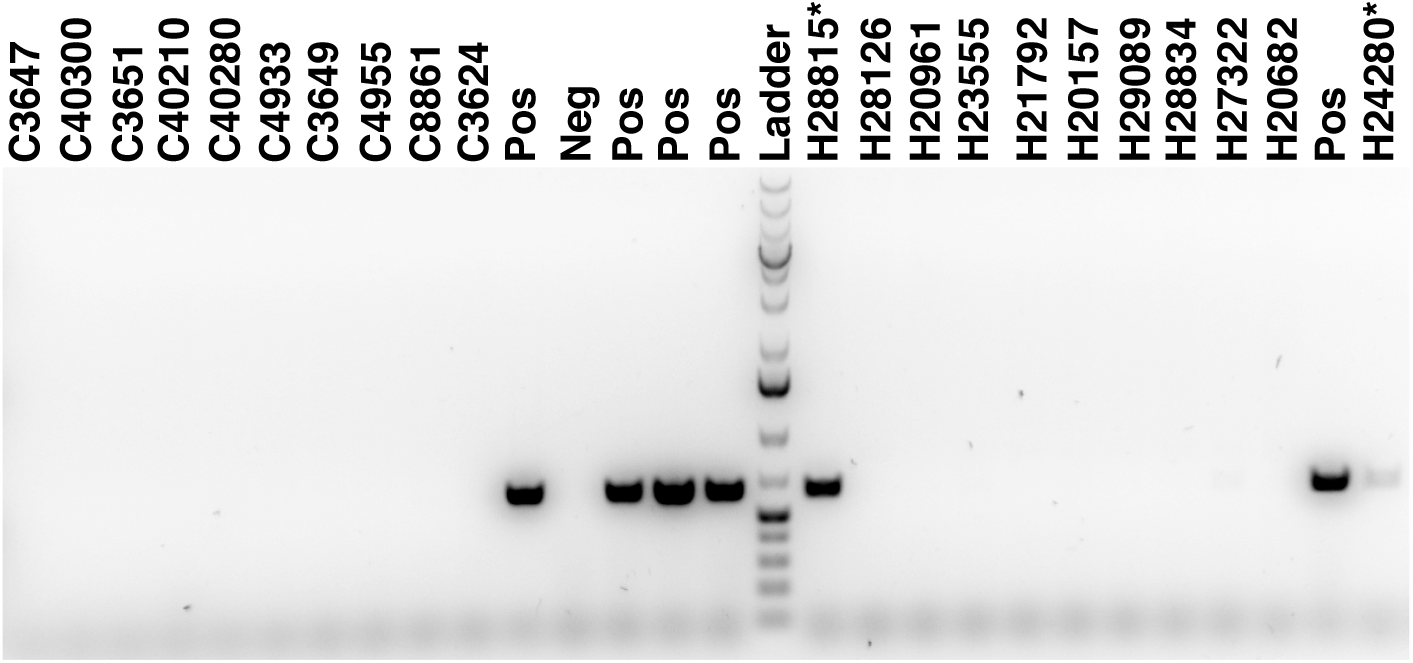
PCR gel to test for exogenous episomal reprogramming vectors. PCR gel for all iPSC lines used for this study. Pos indicates reprograming vector positive DNA controls, Neg is a reprograming vector negative control. Human lines H28815 and H24280 demonstrate positive results for presence of reprogramming plasmid.

**Supplemental Figure 4.**
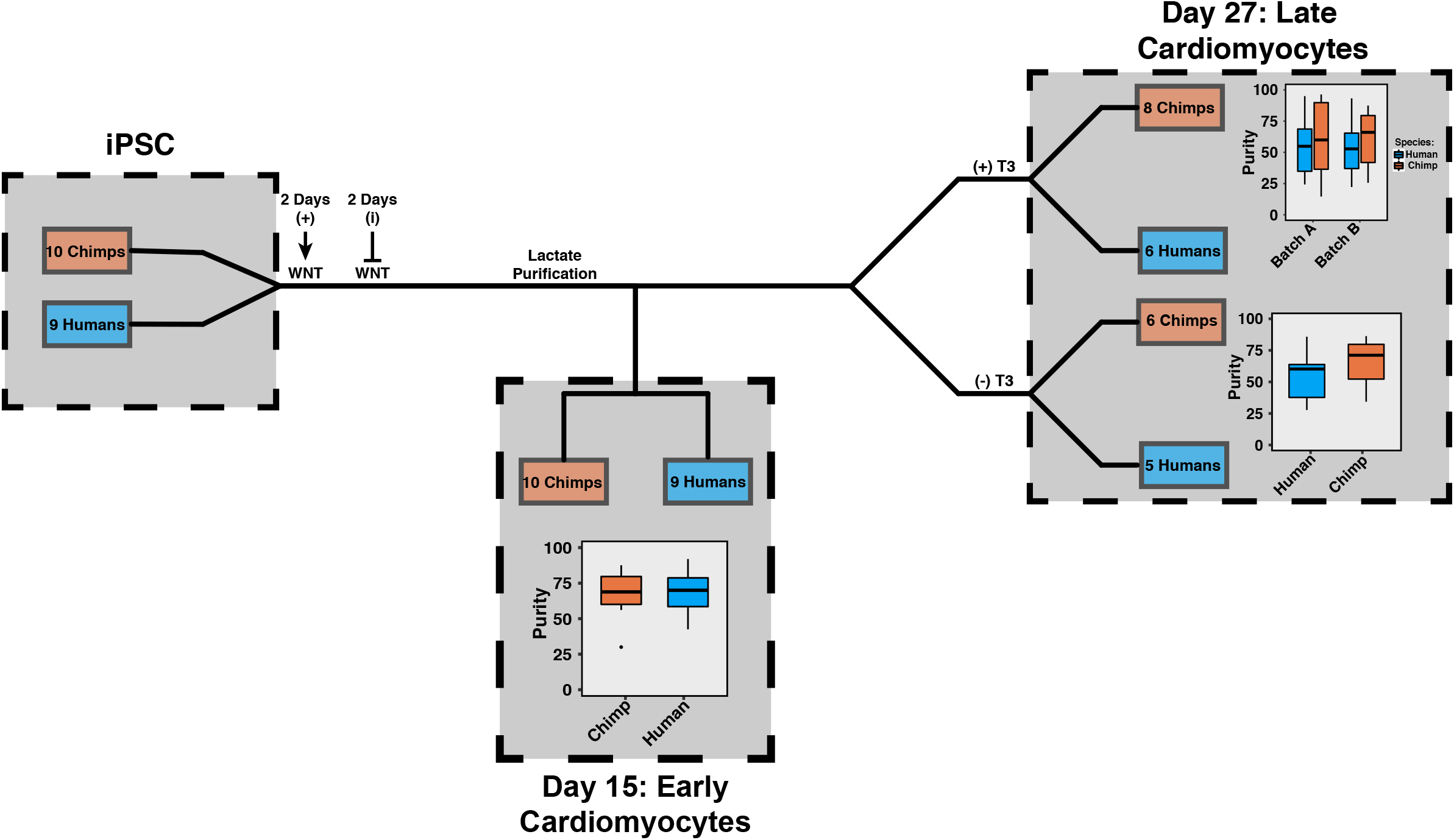
Study design with iPSC-derived cardiomyocytes differentiation outline and summary of data collected. We induced differentiation from iPSC lines using a previously published protocol using WNT agonist for 2 days followed by WNT antagonism for 2 days. At day 15 we harvested RNA from early iPSC-derived cardiomyocytes after metabolic purification with lactic acid. Within a single batch, we collected 10 chimps and 9 human lines. Cells from a subset of individuals were replated and allowed to recover before starting treatment with T3 ((+)T3). A matched set of cells were cultured in parallel but not treated with T3 ((-)T3). We harvested RNA from (+)T3 and (-)T3 day 27 iPSC-derived cardiomyocytes within the same batch that was matched to the day 15 collection (batch B). We also collected another set of (+)T3 treated Day 27 iPSC-derived cardiomyocytes that did not have a matched (-)T3 or Day 15 set (denoted batch A). Purities obtained from dual positive (TNNI3+/TNNT2+) flow cytometry analysis for iPSC-derived cardiomyocytes samples are show in box plots for each set of samples and batches. Grey shaded boxes denote where sample collection occurred.

**Supplemental Figure 5.**
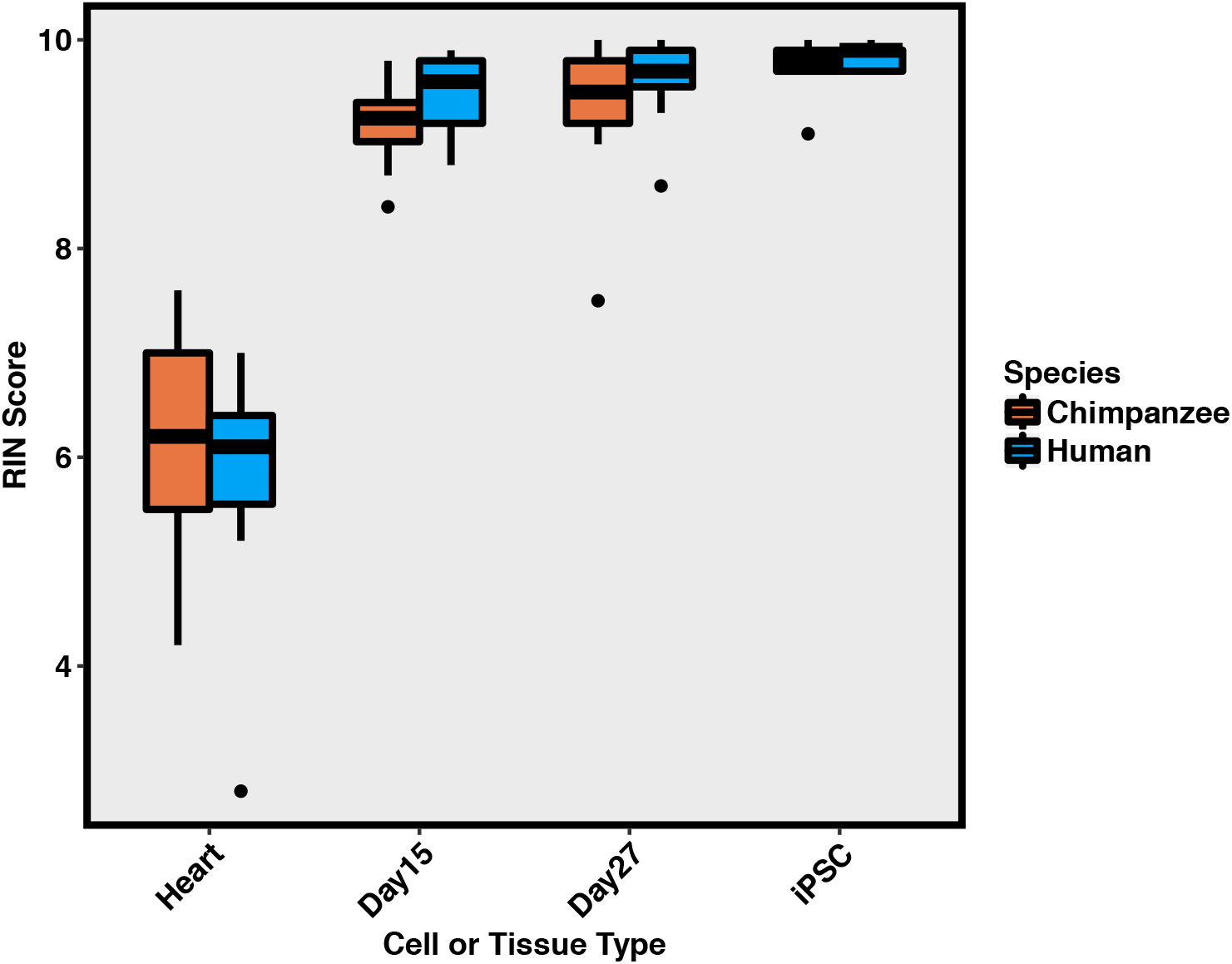
RIN scores for all samples collected in this study. RIN scores separated by species and cell or tissue type for all samples collected in this study.

**Supplemental Figure 6.**
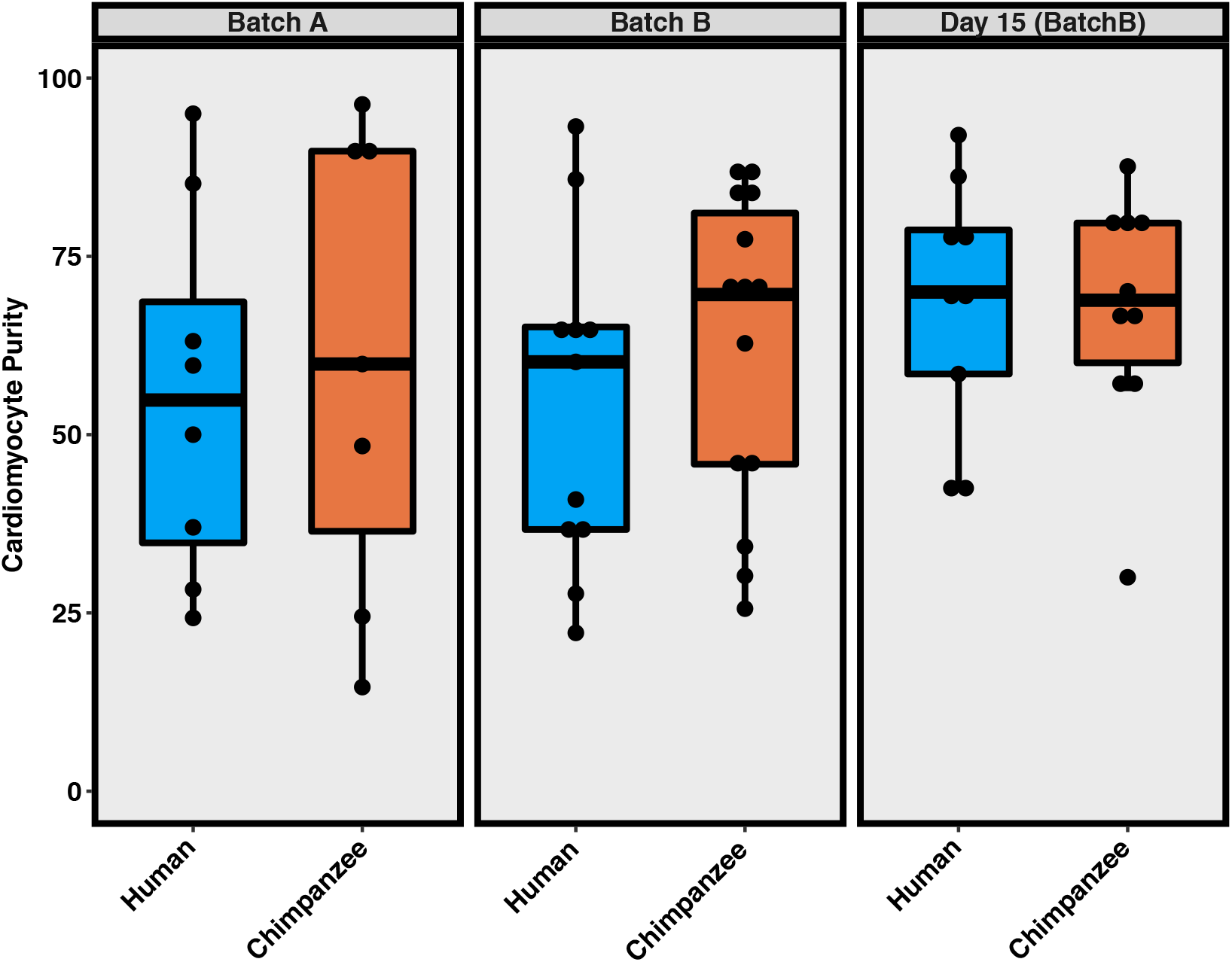
Differentiation batch and purity are not associated with species. Distribution of purities for human and chimpanzee iPSC-derived cardiomyocytes for all samples collected in this study stratified by differentiation batch.

**Supplemental Figure 7.**
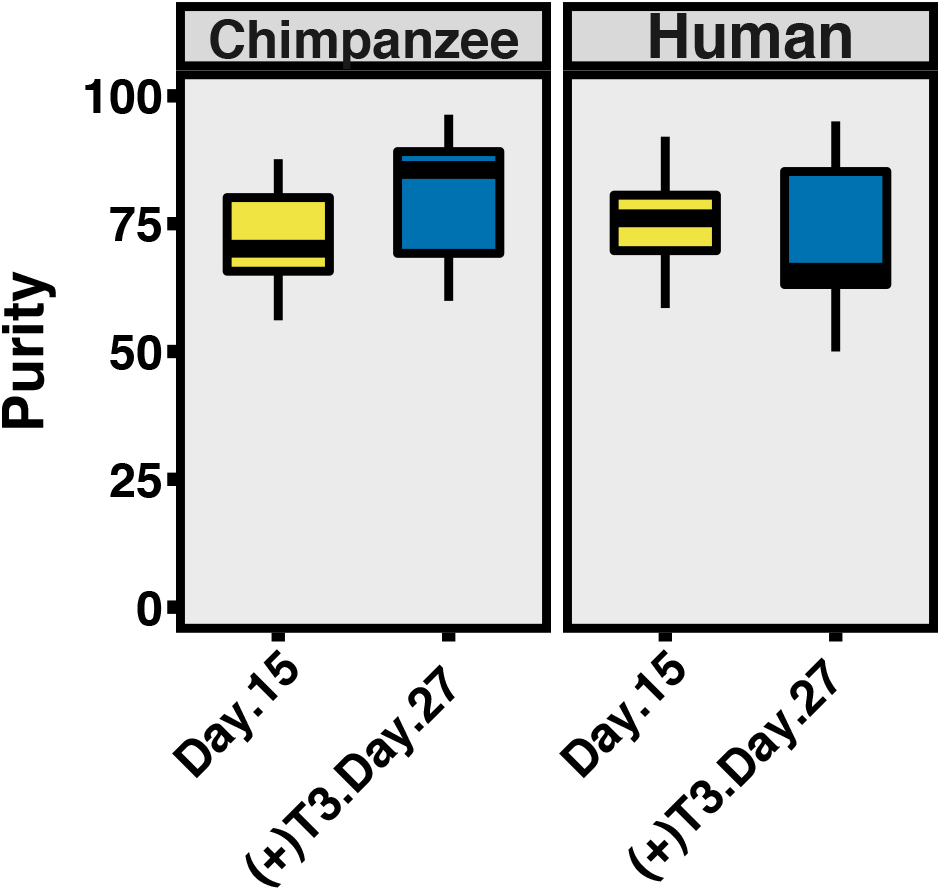
Purity of samples used for analysis of genes that are under differential regulation within iPSCs, iPSC-derived cardiomyocytes and heart tissue. Box plots showing the purity of seven chimpanzee (left) and seven human (right) day 15 and day 27 iPSC-derived cardiomyocytes used for analysis, see Supplemental Table 1 for list of samples used.

**Supplemental Figure 8.**
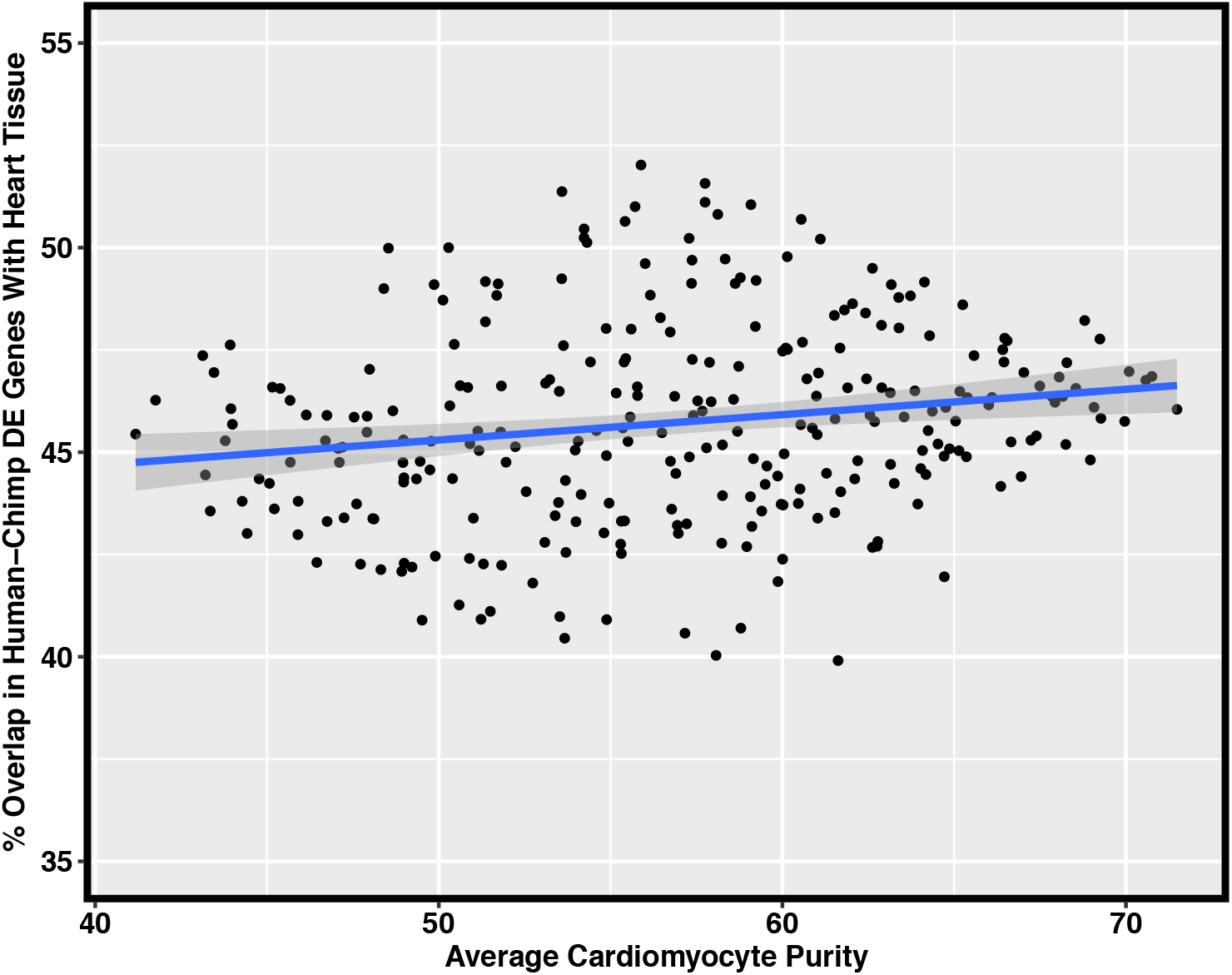
Effect of purity on recapitulation of interspecies DE patterns using iPSC-derived cardiomyocytes. Scatter plot showing the average purity of sample set used on X axis and the percentage of human-chimpanzee DE genes identified on the Y axis. Regression line is shown for Y ∼ X in blue with a 95% confidence interval in gray.

**Supplemental Figure 9.**
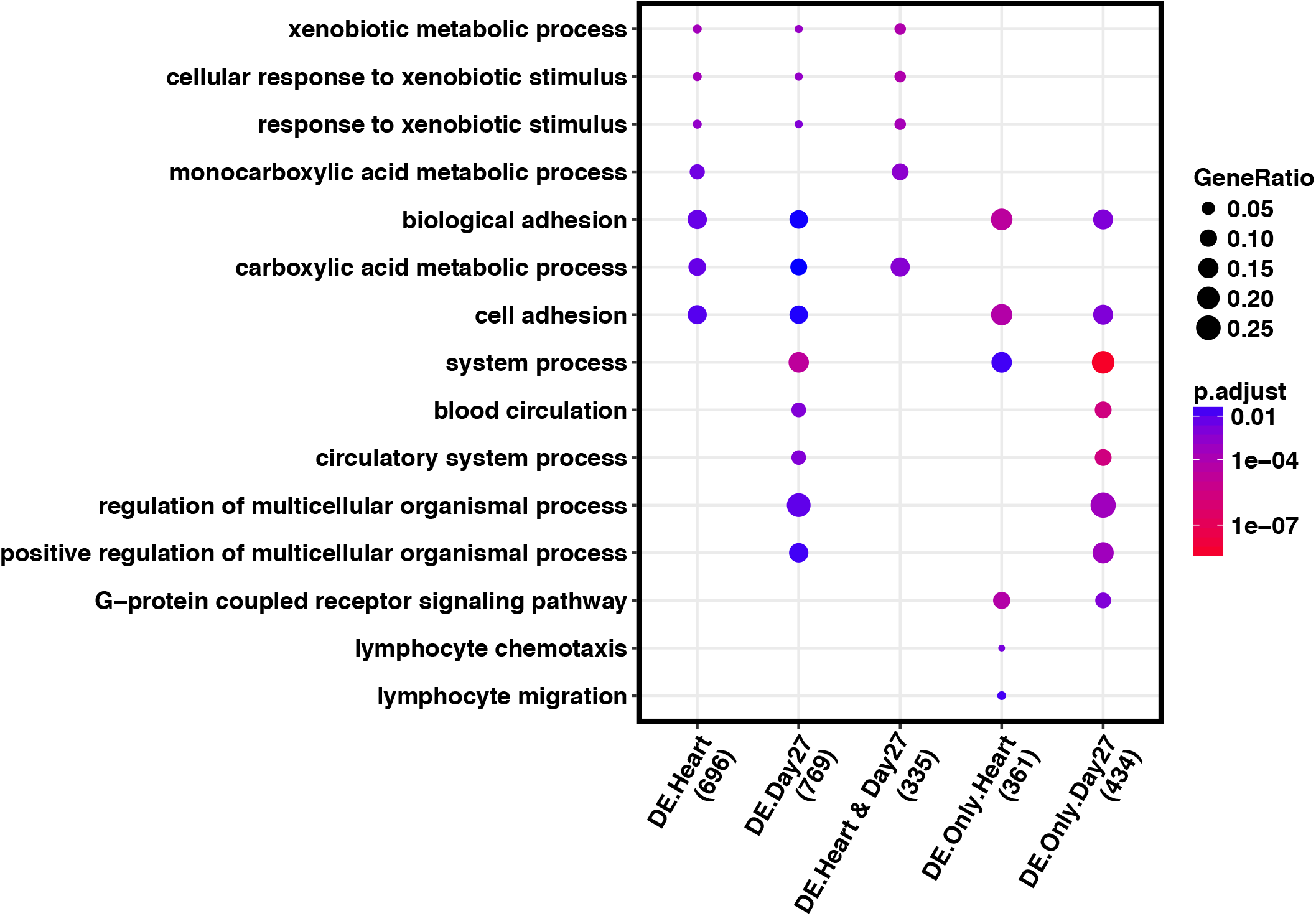
Gene enrichment analysis for interspecies DE genes. The top enrichments in GO biological processes for each of the categories of genes is shown, gene list sizes are shown under the label for each gene set. Complete GO enrichment results are available in Supplemental Table 5.

**Supplemental Figure 10.**
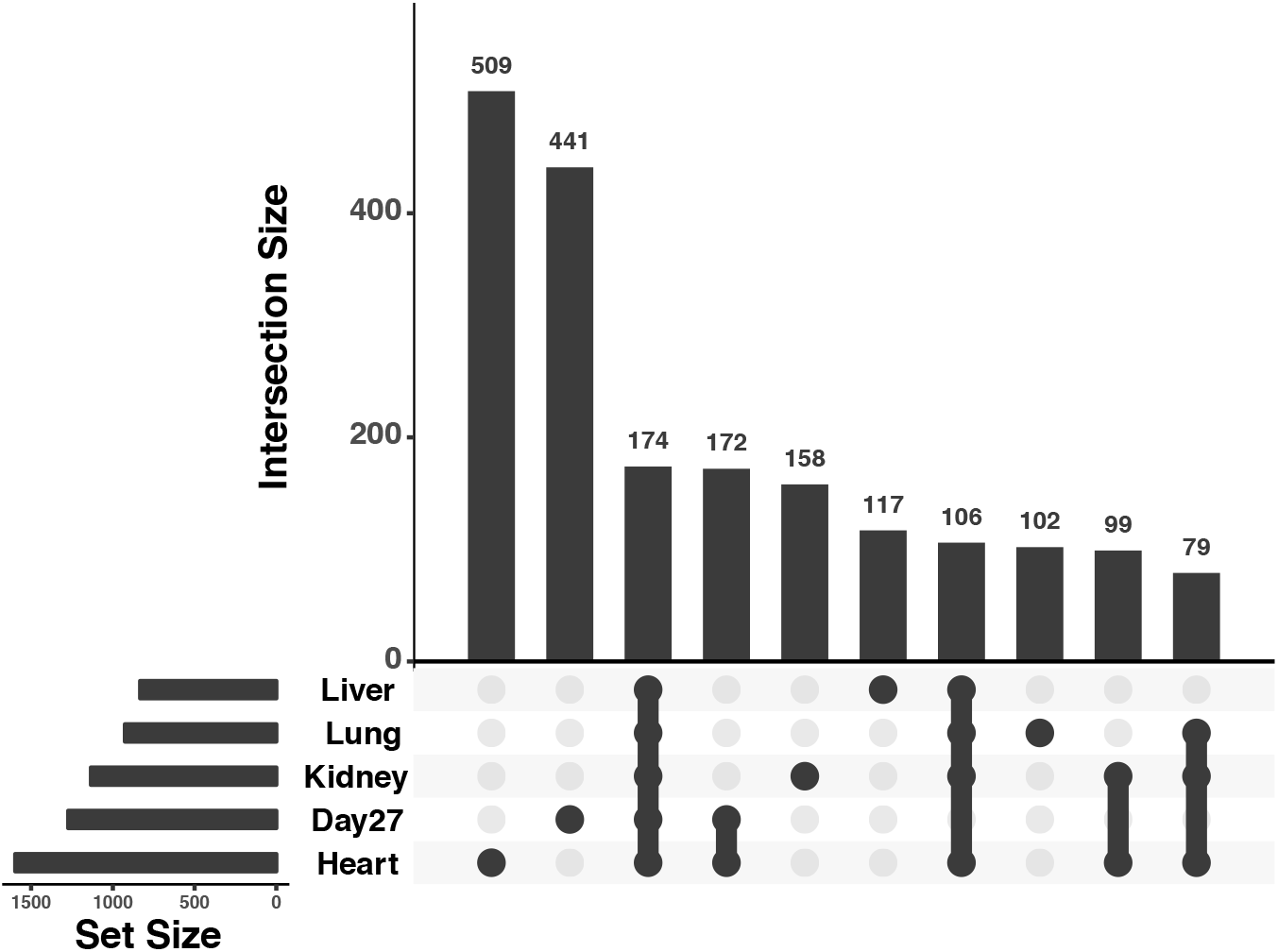
Upsetr plot showing top 10 largest interspecies DE gene overlaps across multiple tissues and day 27 cardiomyocytes. Total set sizes are shown on the bottom left, overlaps are shown by links with a filled circle, and the bar plot above link shows size of a specific overlap. Plot generated using R package Upsetr (Conway et al. 2017).

**Supplemental Figure 11.**
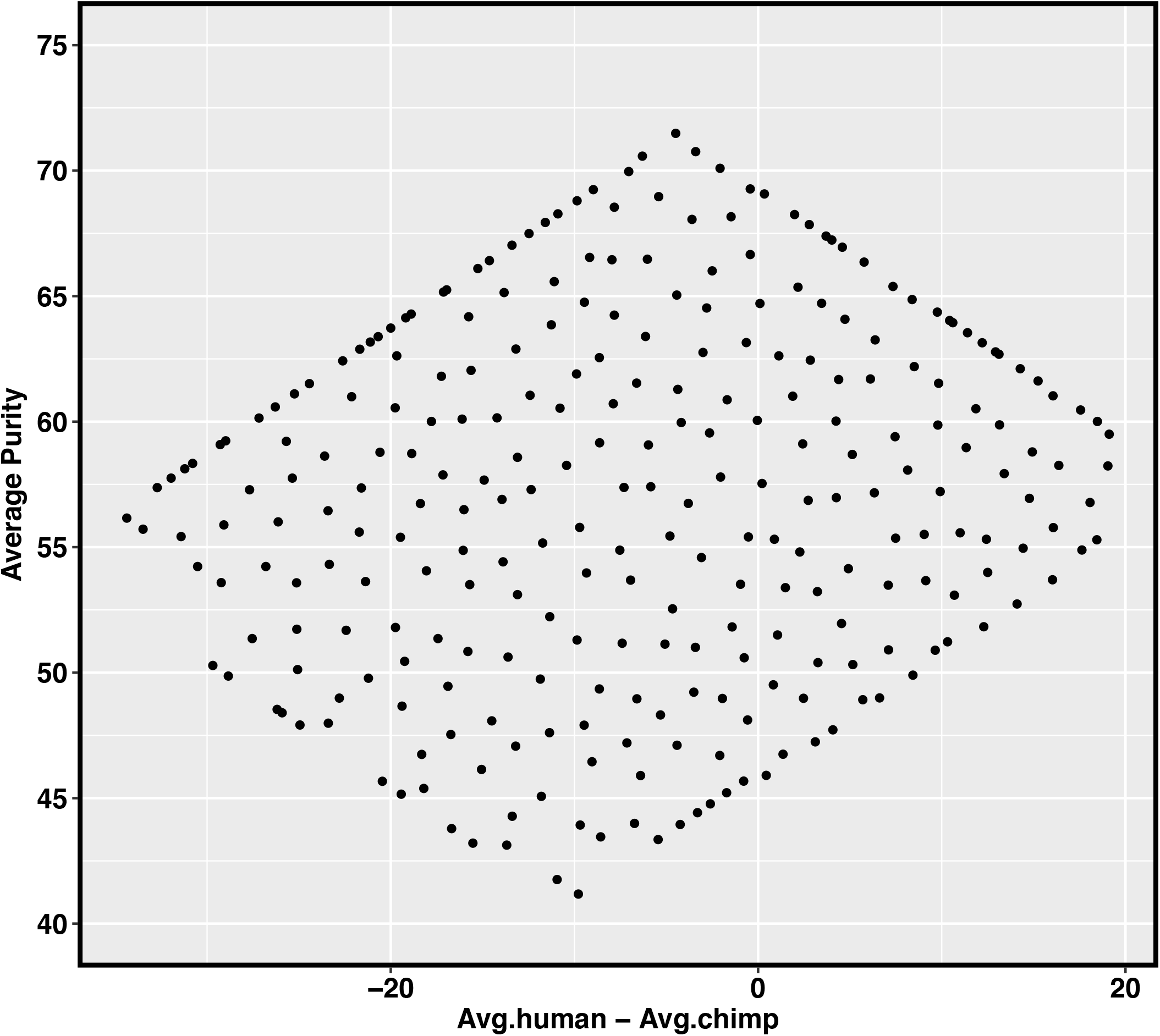
Average purity for all samples and difference in purity between species for samples used to estimate the effect of purity on recapitulating interspecies DE genes. A total of 277 different combinations of average purities for 7 chimps and 7 humans were generated to estimate how purity affects recapitulation of interspecies DE patterns identified in heart tissues. The X axis shows the average purity for all samples together, The Y axis shows the difference in average purity between species.

